# Improvement of protein tertiary and quaternary structure predictions using the ReFOLD4 refinement method and the AlphaFold2 recycling process

**DOI:** 10.1101/2022.12.06.519289

**Authors:** Recep Adiyaman, Nicholas S. Edmunds, Ahmet G. Genc, Shuaa M. A. Alharbi, Liam J. McGuffin

## Abstract

**Motivation:** The accuracy gap between predicted and experimental structures has been significantly reduced following the development of AlphaFold2. However, for further studies, such as drug discovery and protein design, AlphaFold2 structures need to be representative of proteins in solution, yet AlphaFold2 was trained to generate only a few structural conformations rather than a conformational landscape. In previous CASP experiments, MD simulation-based methods have been widely used to improve the accuracy of single 3D models. However, these methods are highly computationally intensive and less applicable for practical use in large-scale applications. Despite this, the refinement concept can still provide a better understanding of conformational dynamics and improve the quality of 3D models at a modest computational cost. Here, our ReFOLD4 pipeline was adopted to provide the conformational landscape of AlphaFold2 predictions while maintaining high model accuracy. In addition, the AlphaFold2 recycling process was utilised to improve 3D models by using them as custom template inputs for tertiary and quaternary structure predictions.

**Results:** According to the Molprobity score, 94% of the generated 3D models by ReFOLD4 were improved. As measured by average change in lDDT, AlphaFold2 recycling showed an improvement rate of 87.5% (using MSAs) and 81.25% (using single sequences) for monomeric AF2 models and 100% (MSA) and 97.8% (single sequence) for monomeric non-AF2 models. By the same measure, the recycling of multimeric models showed an improvement rate of as much as 80% for AF2 models and 94% for non-AF2 models. The AlphaFold2 recycling processes and ReFOLD4 method can be combined very efficiently to provide conformational landscapes at the AlphaFold2-accuracy level, while also significantly improving the global quality of 3D models for both tertiary and quaternary structures, with much less computational complexity than traditional refinement methods.

## 1. INTRODUCTION

There has been a CASP community effort to predict protein structures at high accuracy for three decades (Kryshtafovych et al., 2019; Senior et al., 2019; Subrahamaniam & Kleywegt, 2022). At CASP14, DeepMind’s AlphaFold group submitted tertiary structure models which were widely accepted to represent a step-change in model quality. The models from the AlphaFold2 (AF2) method reached near experimental accuracy for computational modelling. Nevertheless, the actual process of the protein folding path, the effect of conformational flexibility and mutations on functionality, and interacting partners remain unclear (Marx, 2022; Skolnick et al., 2021)

### 1.1 The role of the Molecular Dynamics simulations for improving AF2 structure predictions

Proteins often adopt many different conformations due to their inherent flexibility. In addition to this, different protein conformations could be presented when performing various functions. Although the backbone is well predicted by AF2, it is possible that a model could represent an inactive conformation which does not provide adequate information about its function (Marx, 2022; Skolnick et al., 2021). AF2 predictions were found to be highly accurate in CASP14 (Jumper et al., 2021 (b); Marx, 2022; Skolnick et al., 2021). However, a downside of AF2 is that it provides only a few 3D models rather than conformational dynamics (Jumper et al., 2021 (b); Skolnick et al., 2021), and 3D model comparisons based on mostly C-alpha superpositions may not yield sufficient data to comprehend protein functions. The Molecular Dynamics (MD) protocols, which employ physics-based force fields, may rationalise protein interactions and functions by simulating structural conformational changes that may occur under cellular conditions.

AF2 can predict protein structures at high accuracy while also providing accurate local quality estimates. Protein structure prediction methods have traditionally provided per-residue accuracy scores with the model coordinate data (Varadi et al., 2022). AF2 generates a predicted per-residue accuracy score, which is based on lDDT-Cα and ranges from 0 to 100, with higher scores indicating a better prediction for each residue (Jumper et al., 2021 (b); Marx, 2022; Skolnick et al., 2021).

Since CASP13, we have been pioneering the use of local quality estimates to guide our MD simulation protocols by restraining highly accurate regions and focusing attention on the poorly predicted regions for model refinement. ReFOLD2 was developed by applying a threshold based on the local quality estimation to restrain highly predicted regions in CASP13 (Adiyaman & McGuffin, 2019). In CASP14, the restraint strategy was further developed to apply a gradual restraint rather than a threshold (Adiyaman & McGuffin, 2021). This protocol was made available to the community as the ReFOLD3 web server, which was among the top-performing protocols in CASP14 (Adiyaman & McGuffin, 2021). Here, we have adapted our ReFOLD3 protocol to utilise the AF2 built-in local quality estimates, which are available in the B-factor column in the predicted 3D models. Our aim with ReFOLD4 is to provide conformational changes while maintaining high accuracy in the local regions, and prevent AF2 predictions from structural drift, while using modest computational resources compared to traditional MD-based protocols.

### 1.2 Using the AF2 custom template option with further recycling to improve the quality of input protein structure models

AF2’s algorithmic model is based on two key processes; multiple sequence alignments (MSA) and deep neural networks (DNN), neither of which are especially new or exclusive to AF2. The key to AF2’s recent success appeared to be the construction of detailed residue pair representations used to construct an idealised model to which structures can be repeatedly compared during modelling.

There was also a third interesting process; the existence of a recycling route, which allowed repeated passing of the partially completed proto-model through the DNNs until no further improvement was detectable. If the input of models could be controlled, then this represents a ready-made refinement loop.

ColabFold is a publicly available adaptation of AF2 using a fast MMseqs2 search facility (Mirdita et al., 2022) which also includes a custom template function. This can be adapted to add templates directly into the recycling loop. Our hypothesis for using ColabFold in our CASP15 modelling pathway was that the custom template option could be utilised to input full models into the recycling loop for refinement. Support for this viewpoint comes from the ColabFold team’s recent preprint; State-of-the-Art Estimation of Protein Model Accuracy using AlphaFold (Roney & Ovchinnikov, 2022).

For protein complexes traditional rigid-body docking does not allow for any conformational changes on binding. Thus, refinement becomes an essential step if the model is to be used for drug design or protein-protein interactions (Verburgt & Kihara, 2022). Several docking programs already include flexible modelling, such as iATTRACT (Schindler et al., 2015) and HADDOCK (De Vries et al., 2010), in an attempt to improve natural contacts at the interface and provide more native shape conformity (Schindler et al., 2015).The Deepmind group designed AF2_Multimer (Evans et al., 2021) specifically to model complex structures following Yoshitaka Moriwaki’s work showing that AF2 could be used to model complex protein structures using a linker between two chains.

To test our hypothesis we selected two sets of CASP14 tertiary and quaternary structure models and used the preCASP14-trained AF2 model (see Materials and Methods below) to recycle them. For tertiary structures we chose DeepMind’s AlphaFold group’s (group 427) official CASP14 submissions as well as models selected from the five groups ranking immediately below 427. As AlphaFold did not submit quaternary structures for CASP14, models were generated for the CASP14 targets using AF2-Multimer. Non-AF2 quaternary structure models from the top groups were also used, as described above for monomers.

We attempted to control for the argument that this procedure amounted to simple remodelling in two ways; firstly, by using the official AlphaFold group’s CASP14 models with the AF2 model trained on pre-CASP14 data - the rationale being that the same software should not be able to improve upon its original model. Secondly, by running parallel MSA and single sequence recycling we controlled for any influence that an updated MSA might introduce. Consistent improvement under these conditions would suggest that AF2 is indeed refining the models.

## 2. MATERIALS AND METHODS

### 2.1. The ReFOLD4 protocol

In our ReFOLD4 pipeline, we further improved the MD-based protocol of ReFOLD3 to provide the conformational landscape of the AF2 predictions by applying a unique fine-grained restraint strategy. To fix the local errors identified by the built-in local quality estimation, a fine-grained restraint strategy based on the plDDT score was proposed. In other words, the plDDT score was used as the force constant to be multiplied by the weak harmonic positional restraints (0.05 kcal/mol/Å^2^) (Mirjalili et al., 2014) for each residue on all atoms, including C-alphas, during the MD simulation. As a result, each residue’s restraint sensitivity varies according to its accuracy score, where the restraint ranges from 0.05 kcal/mol/Å^2^ to 5 kcal/mol/Å^2^ instead of determining a local quality estimation score range as in the ReFOLD3 protocol (Adiyaman & McGuffin, 2021).

In our first version of the ReFOLD method (Shuid et al., 2017), the MD simulation parameters were optimized for a modest computational resource compared to other MD-based protocols tested in CASP experiments. The optimised parameters were also used in this MD-based protocol. In summary, the MD simulations were conducted using NAMD version 2.10 via a parallel GPU-based implementation (Phillips et al., 2005). A CHARMM22/27 force field was used to describe the system (Best et al., 2012), the structure was solvated with the TIP3 water model (Jorgensen et al., 1983), and the total charge was neutralized with Na+ or Cl- ions using Particle Mesh Ewald (PME) (Götz et al., 2012). The simulations were performed at 298 K with 1 bar using Langevin dynamics (Loncharich et al., 1992) for temperature and pressure coupling. The default simulation parameters of CHARMM27 were used to exclude non-bonded interactions, which are mostly van der Waal bonds with a switching distance of 10 Å (Mirjalili et al., 2014). Also, hydrogen bonds were rigidified using the rigidBonds function with a 2 fs timestep (Mirjalili et al., 2014). For the first step of MD simulation, an energy minimization protocol for 1,000 steps was applied, followed by the main MD simulation. The fine-grained harmonic positional constraints on all atoms including C-alphas were also performed for 2 ns for each of four parallel simulations for a total of 8 ns. Following the completion of the MD simulation, protein images were generated for each 50ps to generate 164 3D models in total.

LocalColabFold (Mirdita et al., 2022) was used to generate AF2 monomer predictions for CASP14 structures in the default mode (recycle 3) without the template option, and these 3D models were used as starting models for the refinement pipeline. Nevertheless, at the time of running LocalColabFold (March 2022), it was employing the PDB70 database (version Sept. 16, 2020), which was the pre-CASP14 version, so the CASP14 native structures were not seen by LocalColabFold (Mirdita et al., 2022). The best AF2 predictions during CASP14, which were for FM targets, were also further refined using the ReFOLD4 protocol. 16 FM targets were investigated, which had official observed structures available via the CASP website, although for the ReFOLD4 analysis T1096 was removed from ReFOLD4 analysis due to persistent simulation errors.

### 2.2 The AF2 recycling protocol for AlphaFold monomeric models

CASP14 rank 1 AlphaFold tertiary models were downloaded from the CASP website along with their official scores and observed experimental structures. Previous studies have suggested that models created using highly accurate template-based modelling (TBM) have less room for improvement than those created using free modelling (FM) methods (Adiyaman & McGuffin, 2021). Therefore, to maximise the refinement potential and to match the ReFOLD4 process, we used FM models for the16 CASP targets with observed structures.

Two structural alignment scoring methods were used to provide performance metrics for model benchmarking. These scores, generated by downloadable versions of TM-score (Zhang & Skolnick, 2004) and lDDT score (Mariani et al., 2013), describe the backbone (TM-score) and local environment (lDDT) similarities of predicted and observed protein structures. The “baseline” TM and lDDT scores were obtained by comparing the downloaded models for each target from each group with the observed structures.

The model PDB files were then converted to mmCIF format with https://mmcif.pdbj.org/converter using the RSCB PDB MAXIT suite of programs. The converted model files were then submitted to Google Colaboratory hosted ColabFold (release 3, v1.3.0 (4-Mar-2022)) as custom templates along with their respective amino acid sequences. Each model was submitted for 1, 3, 6 and 12 recycles and duplicated for both MSA and single sequence-only modes. ColabFold settings used were: Template_mode: custom; msa_mode: MMseqs2 (UniRef+Environmental) OR single sequence; pair_mode: unpaired+paired; model-type: auto; num_recycles: 1, 3, 6, 12. (N.B. Selecting “auto” from the model type defaulted to monomer_ptm, which was the original pre-CASP14 model).

The five models created for each ColabFold run were collected along with their predicted pTM and plDDT scores. Rank 1 models were then rescored with TM-score and lDDT in the same way as described for baseline scoring. Scores obtained at baseline and for each recycle combination, along with predicted scores (pTM and plDDT), were then directly compared. Statistical analysis was performed using R-studio version 1.3.1093.

### 2.3 The AF2 recycling protocol for non-AlphaFold monomeric models

The same 16 CASP14 targets were selected from the next five best-ranked groups beneath AlphaFold at CASP14. These are (by rank): Baker, Baker-experimental, Feig-R2, Zhang and tFold_human. In addition, to make the test more challenging, only models with a CASP TM-Score of >= 0.45 were used, as those below this threshold cannot be guaranteed to have the same fundamental fold as the reference models (Xu & Zhang, 2010), a total of 47 models were processed.

Models, scores and the observed reference structures for these targets were downloaded from the CASP14 website and scored with TM-score and lDDT in the same way as described in 2.2. ColabFold recycling using MSA was submitted to the same Google Colaboratory version of ColabFold as referenced in 2.2, recycling using single sequence mode was carried out using release v1.3.0 of localcolabfold (Mirdita et al., 2022) installed on our own server, to overcome the Google Colaboratory GPU restrictions in the time available. The equivalent Localcolabfold settings were used: msa-mode: single_sequence; model-type: auto; rank: plddt; pair-mode: unpaired+paired; templates: --custom-template-path.

### 2.4 The AF2 recycling protocol for multimeric models

We repeated the process above but this time using the multimeric CASP14 targets. 10 CASP14 targets were used (H1045, H1065, H1072, T1032, T1054, T1070, T1073, T1078, T1083, T1084) for which the top 5 performing groups had all provided quaternary structure predictions (Baker-experimental, Venclovas, Takeda-Shitaka, Seok & DATE). Since the AF group did not submit any complex target in the CASP14, we also generated AF2-Multimer models for the same targets, so we could then perform common subset analysis.

Each of the baseline models were then refined using the template option and similar parameters as for monomers described above. We also aimed to establish the optimal number of recycles for complex structures. The default value for monomeric AF2 structures is 3 recycles, however, there has been no validation of complex structures, hence we tested out several different numbers of recycles. For scoring the modelled complexes, the MM-Align (Mukherjee & Zhang, 2009) and OpenStructure programs were used for obtaining observed scores for the TM-score and the oligo IDDT score respectively. In addition, the QS-score from OpenStructure was used to simultaneously score interfaces between all interacting subunits.

## 3. Results and Discussion

### 3.1 The performance of the ReFOLD4 pipeline

We used our new ReFOLD4 protocol to refine 57 regular CASP14 targets predicted by LocalColabFold. Out of the 57 targets, 29 were designated as TBM, 15 as FM, and 13 as FM/TBM. The 15 FM best predictions made by the AF2 group (427) during CASP14 were also further refined to test ReFOLD4’s capabilities. Although C-alpha-based assessment scores such as GDT-TS (Zhang & Skolnick, 2004) and lDDT (Mariani et al., 2013) are insufficient to evaluate a structure’s conformational landscape, they were also used alongside the Molprobity score to evaluate ReFOLD4’s performance (Adiyaman & McGuffin, 2019).

In contrast to the GDT-TS and lDDT scores, Molprobity (Chen et al., 2010) takes into account all atoms rather than just C-alpha superpositions as a non-native structure-dependent scoring method. Experimental structures may also be determined at low resolution and may contain some local errors, so the Molprobity score may provide an alternative benchmark by reporting atomic clashes, poor rotamers, and Ramachandran outliers (Chen et al., 2010). It is remarkable to note that on average, 94 percent of the 3D models generated by ReFOLD4 were improved compared to the starting models, and all generated models were improved for 46 targets out of 57 targets according to the Molprobity (see Supplementary Table 1). According to the Molprobity score, ReFOLD4 also significantly outperformed the computationally intensive AF2 group (427) pipeline in CASP14 (∑Molprobitymin = 34.95 and ∑Molprobitymean = 55.86 versus ∑MolprobityAF2CASP14 = 62.91) (Supplementary Table 1).

Our challenge was also to refine the best-submitted models by AF2 group during CASP14 for 15 FM targets to test the fine-grained restraint strategy’s capabilities. It is promising that the fine-grained strategy managed to restrain MD simulations to avoid significant structural deviations according to the GDT-TS and lDDT scores (∑GDTTSmean= 11.35 versus ∑GDTTSstarting= 11.60 and ∑lDDTmean= 10.56 versus ∑lDDTstarting= 11.1) (Supplementary Table 2). A marginal improvement in the overall quality of the best-submitted 3D models was also observed according to the GDT-TS score (∑GDTTSmax= 11.63 versus ∑GDTTSstarting= 11.60) (Supplementary Table 2). ReFOLD4 also showed remarkable success in improving the best-submitted 3D models. Around 72 per cent of the 3D models generated by ReFOLD4 were improved and the cumulative mean and minimum Molprobilty scores were significantly lower than the cumulative Molprobity score of the best-submitted model by AF2 during CASP14 (Table 1).

**Table 1.** Performance summary for ReFOLD4 on the AF2 CASP14 FM models according to Molprobity score

| CASP Target ID | Score of starting model submitted by AF2 in CASP14 | Minimum score of refined models | Mean score of refined models | Maximum score of refined models | Percentage of improved models |
| --- | --- | --- | --- | --- | --- |
| T1027 | 2.39 | 0.83 | 1.217 | 1.6 | 100 |
| T1029 | 0.79 | 0.5 | 0.885 | 1.41 | 40.243 |
| T1031 | 1.62 | 0.5 | 0.968 | 1.42 | 100 |
| T1033 | 1.51 | 0.5 | 0.763 | 1.16 | 100 |
| T1037 | 0.9 | 0.6 | 0.9346 | 1.28 | 100 |
| T1039 | 1.84 | 0.6 | 0.9215 | 1.29 | 100 |
| T1040 | 0.5 | 0.48 | 0.746 | 1.35 | 15.853 |
| T1041 | 1 | 0.52 | 0.889 | 1.24 | 78.658 |
| T1042 | 1.35 | 0.76 | 1.036 | 1.31 | 100 |
| T1043 | 1.2 | 0.51 | 0.950 | 1.42 | 90.243 |
| T1047s1 | 1.61 | 0.73 | 1.095 | 1.49 | 100 |
| T1049 | 0.56 | 0.54 | 0.9535 | 1.55 | 6.097 |
| T1064 | 1.14 | 0.5 | 0.954 | 1.55 | 85.365 |
| T1074 | 1.14 | 0.81 | 1.108 | 1.5 | 63.414 |
| T1090 | 0.68 | 0.52 | 0.932 | 1.38 | 6.097 |
| The Cumulative scores | 18.23 | 8.9 | 14.357 | 20.95 |  |

It is assumed that the ReFOLD4 protocol may perform better on models for FM targets, where there is often more room for improvement. ReFOLD4 was able to produce a larger population (> 10%) of improved models for 7 out of the 15 targets for the refinement of the starting models generated by LocalColabFold, according to the GDT-TS score (Supplementary Table 3). This indicates that ReFOLD4 can generate a sizable portion of improved models for FM targets. The cumulative maximum GDT-TS score was slightly higher than the cumulative GDT-TS score of the starting models (∑GDT-TSmax = 10.80 versus ∑GDT-TSstarting = 10.75) while the cumulative mean GDT-TS score was slightly lower than the cumulative GDT-TS score of the starting models (∑GDT-TSstarting = 10.75 versus ∑GDT-TSmean = 10.56) (Supplementary Table 3). The higher maximum cumulative GDT-TS score means that ReFOLD4 improved the starting models. A similar trend was observed for the FM/TBM targets.

Historically, the refinement of TBM targets has been more challenging, and their refinement is likely to deteriorate the global quality as there is much less room for improvement (Adiyaman & McGuffin, 2019). The role of the restraint strategy also becomes more apparent in TBM targets as unrestrained MD simulations are likely to substantially deviate from the native basin (Adiyaman & McGuffin, 2019, 2021). Although there is much less improvement in the overall quality of the 3D models (∑GDT-TSmax = 23.62 versus ∑GDT-TSstarting = 23.49) on 29 TBM targets, the fine-grained restraint strategy managed to direct the generation of 3D models towards the native basin according to the GDT-TS score (∑GDT-TSstarting = 23.49 versus ∑GDT-TSmean = 23.17) as in Supplementary Table 3. It is worth noting that the majority of 3D models generated by ReFOLD4 were improved for T1060s3, T1092, T1093, T1094 and T1095 (Figure 1, and Supplementary Table 3).

**Figure 1.**
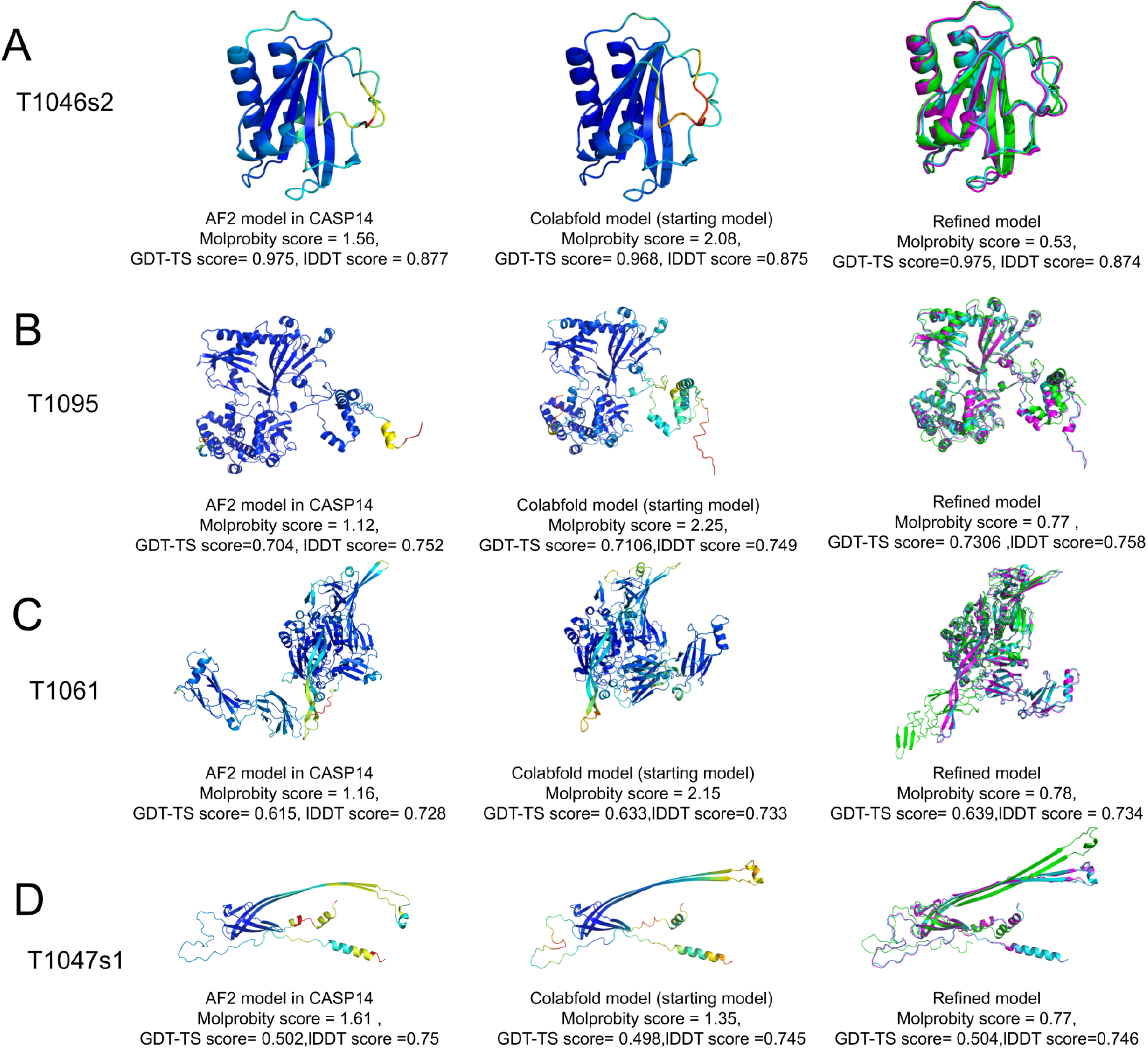
The refinement of four example CASP14 targets using the ReFOLD4 protocol. The images in the left columns show the best models submitted by the AF2 group (427) in CASP14 and were coloured by the plDDT. The middle columns show the best models generated by LocalColabFold coloured by the plDDT, and these models were used as initial structures by ReFOLD4. The right columns show the superposition of the best 3D models generated by LocalColabFold (cyan), the best 3D models generated by ReFOLD4 (magenta) and native structures (green). (A) CASP14 regular TBM T1046s2: the AF2 group model has a Molprobity score of 1.56, a GDT-TS score of 0.975, and a lDDT score of 0.877, the LocalColabfold model was refined using ReFOLD4, with a Molprobity improvement from 2.08 to 0.53 (lower Molprobity scores are better), and a GDT-TS improvement from 0.968 to 0.975. (B) CASP14 regular TBM T1095: the AF2 group model has a Molprobity score of 1.12, a GDT-TS score of 0.704 and a lDDT score of 0.752, the LocalColabfold model was refined using ReFOLD4, with a Molprobity improvement from 2.25 to 0.77, and a GDT-TS improvement from 0.7106 to 0.7306. (C) CASP14 regular FM/TBM T1061: the AF2 group model has a Molprobity score of 1.16, a GDT-TS score of 0.615, and a lDDT score of 0.728, the LocalColabfold model was refined using ReFOLD4, with a Molprobity improvement from 2.15 to 0.78, and a GDT-TS improvement from 0.633 to 0.639. (D) CASP14 regular FM T1047s1: the AF2 group model has a Molprobity score of 1.61, a GDT-TS score of 0.502, and a lDDT score of 0.75, the LocalColabfold model was refined using ReFOLD4, with a Molprobity improvement from 1.35 to 0.77, and a GDT-TS improvement from 0.498 to 0.504. Images were rendered using PyMOL.

The ReFOLD4 pipeline, which used models generated by LocalColabfold as starting models, provided better predictions for 18 targets out of 57 targets (T1030, T1034, T1039, T1046s2, T1047s1, T1047s2, T1049, T1053, T1055, T1061, T1068, T1070, T1078, T1082, T1083, T1084, T1089, and T1095) compared to the AF2 group’s (427) best submissions according to the maximum GDT-TS score (Supplementary Table 3).

The lDDT score (Mariani et al., 2013) which is a native structure-dependent and C-alpha-based score, was also used to analyse the performance of the ReFOLD4 pipeline. Based on this score, ReFOLD4 did not manage to improve most of the targets (Supplementary Table 4). The AF2 method is extremely well-trained to optimise the lDDT score, which is the main target measurement for the training dataset (Jumper et al., 2021 (b); Skolnick et al., 2021). Therefore, further improvements in the overall lDDT quality might not be as achievable by MD simulations. Nevertheless, the fine-grained restraint strategy demonstrated exceptional performance for T1092 and T1095 according to the population of the improved models (Supplementary Table 4).

### 3.2 Performance of the AF2 recycling protocol for refining monomers

As measured by average change in lDDT, monomeric recycling showed an improvement rate of 87.5% (using MSAs) and 81.25% (using single sequences) for AF2 models and 100% (MSA) and 97.8% (single sequence) for non-AF2 models.

A comparison matrix of the 16 CASP14 AF2 and 47 non-AF tertiary structure models is presented in Table 2, below. Calculated p-values were computed from a 1-tailed Wilcoxon signed-rank test for lDDT scores (Table 2A) between the baseline values and the output models from each recycle, and also for output models between consecutive recycles. Identical analyses are presented for TM-score in Table 2B. A significant difference between any two populations is established by a p-value of ≤0.05 suggesting an improvement in quality at the named recycle.

**Table 2.** Calculated p-values, based on lDDT scores (A) and TM-scores (B), for recycled models for CASP14 AF2 and non-AF models for monomeric targets

| A |  |  |  |  |  |  |  |  |
| --- | --- | --- | --- | --- | --- | --- | --- | --- |
| Models | Recycle type | Baseline to 1 recycle | 1 recycle to 3 recycles | Baseline to 3 recycles | 3 recycles to 6 recycles | Baseline to 6 recycles | 6 recycles to 12 recycles | Baseline to 12 recycles. |
| AF2 | MSA | 1.87e-01 | 7.56e-01 | **5.66e-03** | 4.36e-02 | **7.72e-03** | 3.51e-01 | **1.30e-02** |
|  | SS | **1.12e-02** | 9.54e-01 | **1.86e-02** | 1.24e-01 | 5.90e-02 | 6.37e-01 | **3.86e-02** |
| non-AF | MSA | **1.23e-09** | **1.21e-08** | **7.10e-15** | **1.56e-02** | **7.10e-15** | 4.73e-01 | **1.23e-09** |
|  | SS | **1.70e-09** | **4.91e-05** | **1.50e-09** | 1.75e-01 | **1.50e-09** | 5.87e-01 | **1.40e-09** |
| B |  |  |  |  |  |  |  |  |
| Models | Recycle type | Baseline to 1 recycle | 1 recycle to 3 recycles | Baseline to 3 recycles | 3 recycles to 6 recycles | Baseline to 6 recycles | 6 recycles to 12 recycles | Baseline to 12 recycles. |
| AF2 | MSA | 6.79e-01 | 8.01e-01 | 7.96e-01 | 1.06e-01 | 9.58e-01 | 3.63e-01 | 7.76e-01 |
|  | SS | 7.17e-01 | 9.09e-01 | 8.60e-01 | **3.37e-02** | 8.97e-01 | 7.82e-01 | 6.98e-01 |
| non-AF | MSA | **1.42e-14** | **6.31e-05** | **4.36e-12** | 8.98e-01 | **8.05e-09** | 2.40e-01 | **2.40e-12** |
|  | SS | **5.13e-07** | **7.61e-05** | **3.35e-07** | **3.30e-02** | **1.98e-07** | 6.60e-01 | **1.62e-07** |

\*SS=Single sequence. Ho: Recycling as specified per column produces models that are no different in quality to those input as baseline templates or to the previous recycle number. H1: Recycling as specified per column produces higher quality
models to those input at baseline or to the previous recycle number. P-values $\leq 0.05$ indicate significant differences (in bold) were calculated by a 1-tailed Wilcoxon signed-rank test using IDDT score (A) and TM-scores (B) for 16 AlphaFold2 CASP14 top-ranked models (AF2) and 47 non-AlphaFold models (Non-AF) from the same CASP14 targets.

For lDDT score, given the 0.05 p-value threshold, it can be concluded that recycling significantly improved AF2 model quality compared to the baseline models using MSA and single sequence inputs (values shown in bold), although the improvement in model quality was non-linear; higher recycle numbers did not always show greater improvement.

Further, recycling significantly improved non-AF model quality compared to baseline across all recycles for both MSA and single sequence methods. Improvement in model quality again appeared to be non-linear; recycle 6 to 12 produced no further significant improvement for either method.

Identification of the optimal recycle number was not immediately obvious from the data. However, considering the minimal improvement from recycle 3 onwards for non-AF models and the increasing magnitude of the calculated p-values from 3 recycles onwards for AF2 models, it is reasonable to conclude that 3 recycles represented improvement at least as good as any other recycle. Therefore, for this population of monomers we will describe recycle 3 as the optimal number of recycles for refinement of monomeric models.

Figure 2 shows the change in individual model quality for lDDT (Figure 2A) and TM-score (Figure 2B) for all models (MSA recycling) with points coloured by group. A comparison of the plots for 3 recycles and all recycles is shown in Supplementary Figure 1.

**Figure 2.**
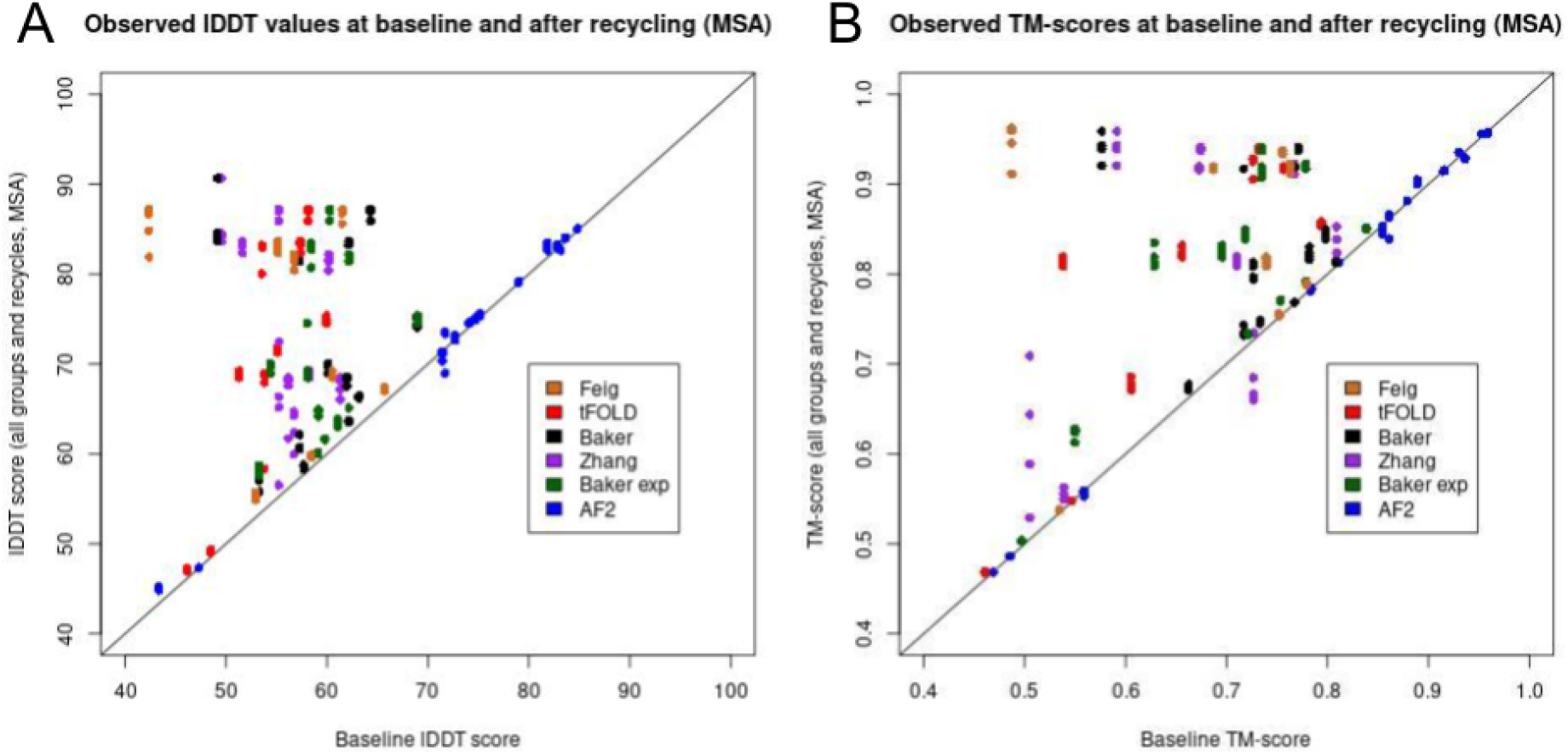
Scatter plots showing the comparison in observed lDDT scores (A) and observed TM-scores (B) between baseline (x-axis) and all recycles (y-axis) for all AF2 and non-AF models (MSA recycling), coloured by group.

Considering lDDT scores, Figure 2A shows that the improvement of AF2 models (blue points), although significant, was relatively minor in real terms. This is, perhaps, unsurprising considering the high baseline lDDT scores associated with the AF2 template models and in fact, expected, considering that refinement is associated with small improvements in atomic positions of an already representative model. As explained in the introduction, it is assumed unlikely that any significant remodelling occurred when using official AlphaFold CASP14 structures coupled with the pre-CASP14-trained AF2 model, as the same software should not be able to improve upon its original model without additional information. It is therefore likely that this improvement represented true refinement by the AF2 algorithm.

For both panels, Figure 2 shows a more noticeable improvement for non-AF models and this is potentially explained by two factors. Firstly, by the lower starting quality of the baseline template models providing greater potential for improvement but, secondly, the possibility that a certain amount of remodelling was taking place during the recycling. It is therefore necessary to identify the extent of this re-modelling and attempt to demonstrate that significant improvement still occurred by recycling alone. To do this we compared single-sequence directly with MSA recycling.

### 3.3 Improvement in MSA versus single sequence recycling

The 1-tailed Wilcoxon signed-rank test was again used to calculate p-values between the two model populations. In addition, a 1-tailed Ansari-Bradley test was used to investigate any significant differences in quartiles to rule out differences occurring in the data unrelated to the mean (p-values are available in Supplementary Table 5).

It was found that there was no significant difference in model quality between MSA and single sequence recycling for the AF2 models according to both the Wilcoxon and Ansari tests. However, there was a significant difference, detected by both tests, at every recycle for non-AF models. This supports the conjecture in 3.1 above, that no significant remodelling occurred for AF2 models but that a certain amount probably occurred for non-AF models with MSA recycling. The difference between the two graphs may represent the additional remodelling afforded by the MSA for non-AF groups (shown in an equivalent plot in Supplementary Figure 2).

Actual changes in lDDT are represented as a bar charts in Figure 3A and Supplementary Figure 3. Figure 3A shows the change in cumulative lDDT for all groups across all MSA recycles and the changes in TM-scores are shown in 3B.

**Figure 3.**
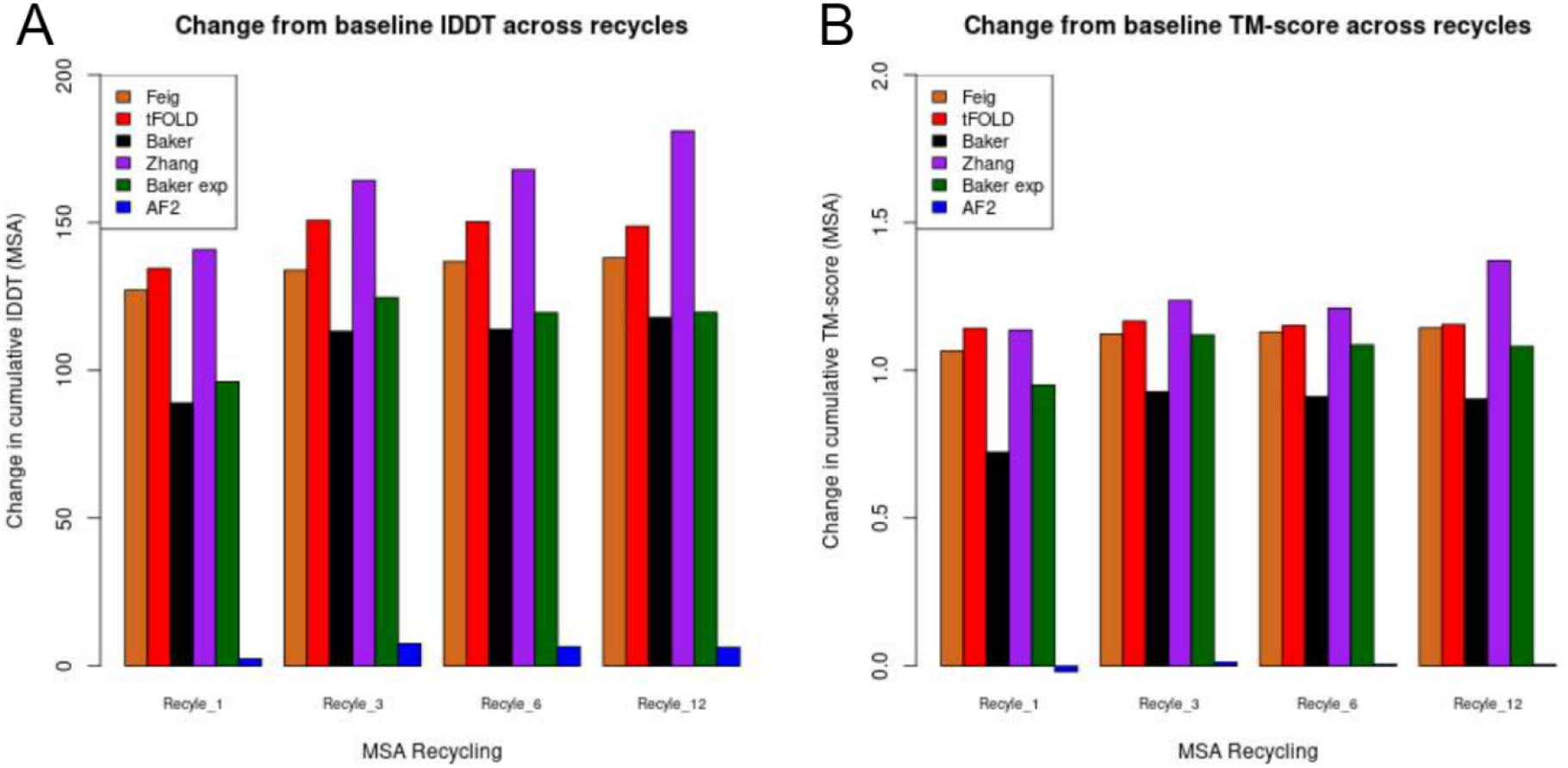
Bar charts showing the cumulative increase in observed lDDT scores (A) and TM-scores (B) from the baseline models to the models produced using different numbers of recycles. Bars are coloured by group.

Both panels clearly show the differences in improvement in model quality obtained between AF2 (blue) and non-AF models (brown, red, black, purple and green columns), again highlighting the differential effect MSA has on the two groups of models. An additional important observation was that many non-AF models with relatively low initial lDDT scores were refined to a level that out-performed both the initial and the refined equivalent AF2 models. Examples of these are shown in Figure 4 below.

**Figure 4.**
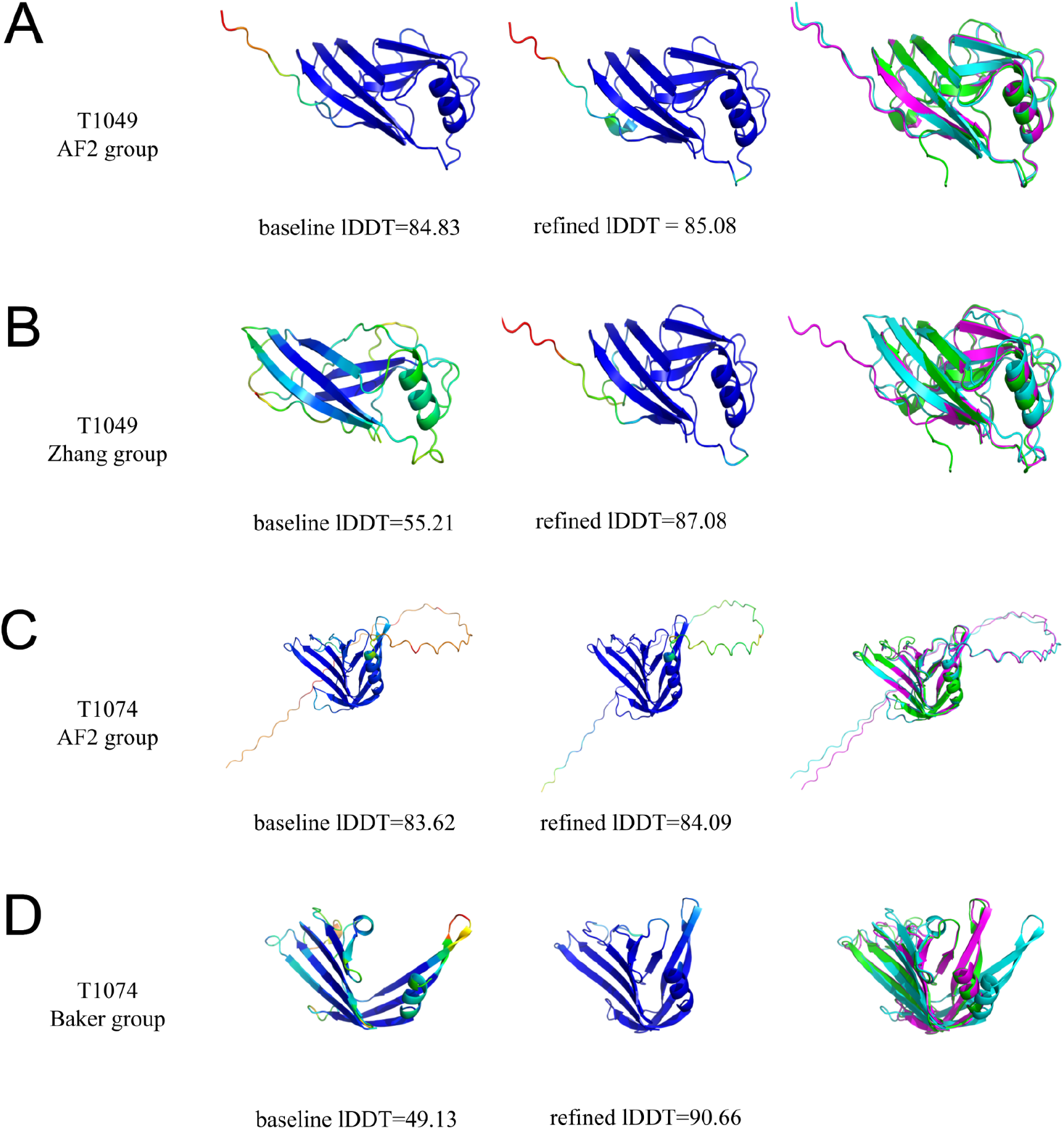
Comparison of monomer models. Images in the left columns show the baseline models coloured by plDDT score. The middle columns show the refined models coloured by plDDT score. The right columns show the superposition of the baseline models (cyan), the refined models generated by colabfold (magenta) and the observed native structures (green). (A) AF2 model for T1049: baseline lDDT=84.83, refined lDDT = 85.08. (B) Zhang group model for T1049: baseline lDDT=55.21, refined lDDT=87.08. (C) AF2 model for T1074: baseline lDDT=83.62, refined lDDT=84.09. (D) Baker group model for T1074: baseline lDDT=49.13, refined lDDT=90.66. Images were rendered using PyMOL.

### 3.4 Performance of the AF2 recycling protocol for refining multimers

According to lDDT scores, recycling of multimer models showed an improvement rate of 80% (using MSAs) and 30% (using single sequences) for AF2 models, while 94% (MSA) and 64% (single sequence) non-AF2 models were improved. According to TM-scores, there was an improvement of 70% (MSA) and 80% (single sequence) for AF2 multimeric models, and 98% (MSA) and 82% (single sequence) for non-AF2 multimeric models. While, according to QS scores, 50% (MSA) and 30% (single sequence) of AF2 multimeric models, and 86% (MSA) and 60% (single sequence) of non-AF2 multimeric models were improved.

While the principal improvement from recycling monomers is evidently the significant gain in lDDT scores, this is not always the case for multimeric models. When we use the same methodology for recycling complexes, the larger improvements are shown to the TM scores and interface quality scores (QS-scores) than for the oligo-lDDT scores (Table 3). That said, compared to the baseline non-AF models, significant improvements were made for all numbers of recycles for all scores when MSAs were used. Furthermore, significant improvements to the both TM-score and QS-scores for non-AF2 models were seen for all recycles when only a single sequence was used. The AF2M models were more consistently significantly improved in terms of their TM-scores, rather than the other scores. Considering the p-values for all models and scores, both 6 and 12 recycles appear to be optimal parameters for significantly improving upon baseline models.

**Table 3.** Calculated p-values based on oligo-lDDT scores (A), TM scores (B), QS-scores (C), for baseline and recycled models for both CASP14 non-AF2 models and the AF2-Multimer models for multimeric targets.

| A |  |  |  |  |  |  |  |  |
| --- | --- | --- | --- | --- | --- | --- | --- | --- |
| Models | Recycle type | Baseline to 1 recycle | 1 recycle to 3 recycles | Baseline to 3 recycles | 3 recycles to 6 recycles | Baseline to 6 recycles | 6 recycles to 12 recycles | Baseline to 12 recycles |
| AF2M | MSA | 1.106e-1 | 5.203e-1 | 1.795e-1 | 7.679e-2 | **4.157e-2** | 1.106e-1 | **5.146e-2** |
|  | SS | 9.966e-1 | **4.157e-2** | 9.369e-1 | 6.177e-2 | 9.369e-1 | 9.736e-1 | 9.369e-1 |
| non-AF | MSA | **3.748e-3** | **4.267e-05** | **1.398e-05** | **6.915e-3** | **1.02e-06** | 9.565e-1 | **4.935e-07** |
|  | SS | 8.492e-1 | **1.475e-2** | 5.116e-1 | 1.611e-1 | 4.197e-1 | **1.013e-2** | 3.285e-1 |
| B |  |  |  |  |  |  |  |  |
| AF2M | MSA | **3.327e-2** | 2.704e-1 | 6.314e-2 | 6.314e-2 | **2.075e-2** | 8.205e-1 | 1.54e-1 |
|  | SS | **5.146e-2** | 1.795e-1 | **2.075e-2** | 6.201e-1 | **1.247e-2** | 6.202e-1 | **2.075e-2** |
| non-AF | MSA | **2.066e-09** | 2.715e-1 | **1.453e-09** | 9.428e-1 | **2.926e-09** | 9.858e-1 | **6.889e-09** |
|  | SS | **3.338e-3** | **3.97e-3** | **5.516e-4** | 7.458e-1 | **1.367e-4** | 3.75e-1 | **2.946e-4** |
| C |  |  |  |  |  |  |  |  |
| AF2M | MSA | 4.161e-1 | 5.724e-1 | 1.976e-1 | 4.27e-1 | 5e-1 | 5e-1 | 5e-1 |
|  | SS | 7.992e-1 | **5.017e-2** | 5e-1 | 1.855e-1 | 3.422e-1 | 8.618e-1 | 3.375e-1 |
| non-AF | MSA | **1.577e-07** | 2.268e-1 | **2.578e-07** | 1.575e-1 | **1.09e-07** | 2.326e-1 | **6.799e-08** |
|  | SS | **3.491e-2** | **1.118e-2** | **4.175e-3** | 3.083e-1 | **2.548e-3** | 2.406e-1 | **4.089e-3** |

\*SS=Single sequence. Ho: Recycling by the number specified per column produces models that are equal or lower in quality than those input as baseline templates or by the previous recycle number. H1: Recycling by the number specified per column produces higher quality models than those input at baseline or by the previous recycle number. P-values ≤0.05 indicate significant differences (in bold). The 1-tailed Wilcoxon signed-rank sum test P-values were calculated using IDDT scores (A), TM-scores (B), QS-scores (C) across 10 AlphaFold models (10 AlphaFold generated by MSA for the MSA method and 10 AlphaFold generated by Single sequence for the Single sequence method) and 50 non-AlphaFold models from different CASP14 targets.

The plots in Figure 5 show that the majority models are improved according to all scoring methods, but again it is clear that the improvement is less consistent for the oligo-lDDT scores (Figure 5A) than for the TM-scores (Figure 5B) and QS scores (Figure 5C). In Figure 5B, the improvement of TM score for both AF2 and non-AF2 is clearer, and far fewer targets were degraded following increases in the recycling number. Similar plots are shown for each recycle separately in Supplementary Figure 4 and plots for single sequence inputs in Supplementary Figure 5 and 6.

**Figure 5.**
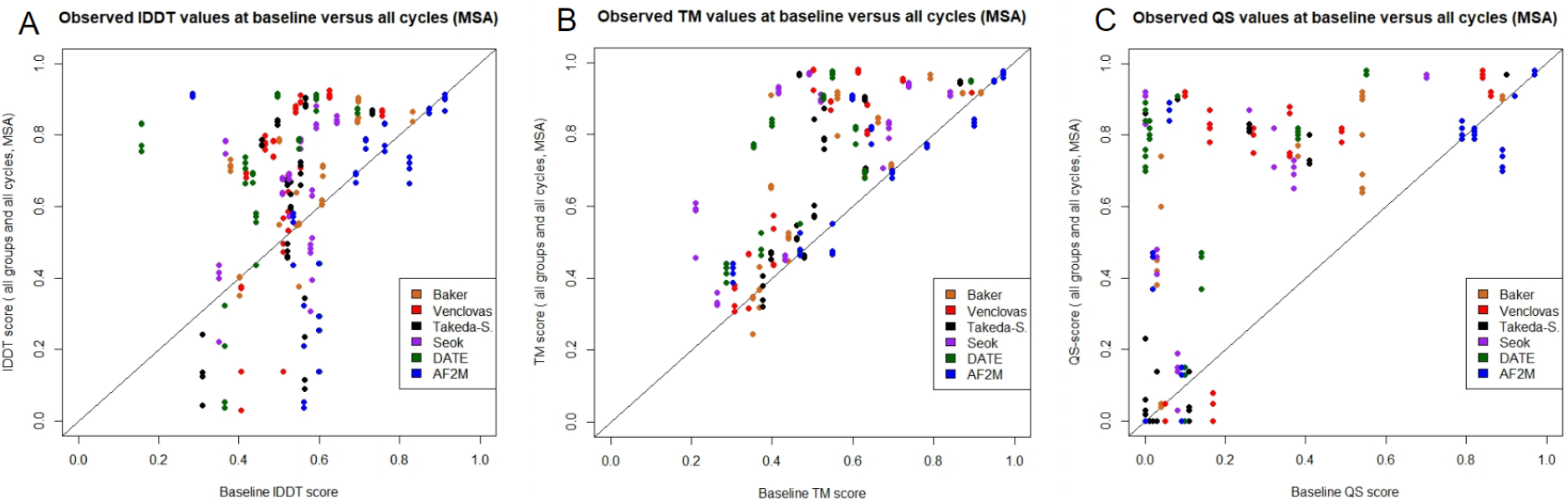
Scatter plots showing the comparison in observed oligo-lDDT (A), observed TM (B), and observed QS scores (C) between baseline (x-axis) and all recycles (y-axis) for all AF2M and non-AF2 multimer models.

Figure 6 shows that the cumulative improvement in model quality was non-linear with increasing numbers of recycles; higher recycle numbers (>3) did not necessarily always lead to greater improvement for all model types according to all scores. However, with 6-12 recycles models from almost all groups can be improved, with clear cumulative gains shown for each score.

**Figure 6.**
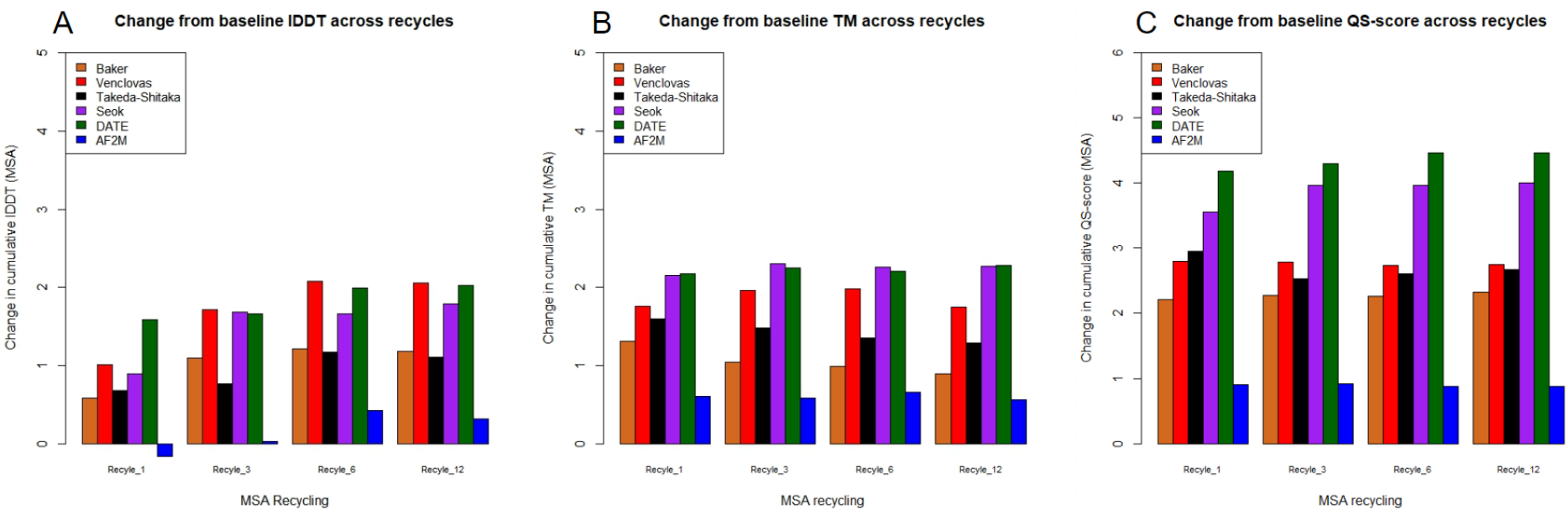
Bar charts showing the cumulative increase in observed oligo-lDDT scores (A), TM-scores (B) and QS score (C) from the baseline multimer models to the models produced using different numbers of recycles. Bars are coloured by group.

Figure 7 shows examples of the visible improvements to the quality of multimeric models from different groups following the recycling process.

**Figure 7.**
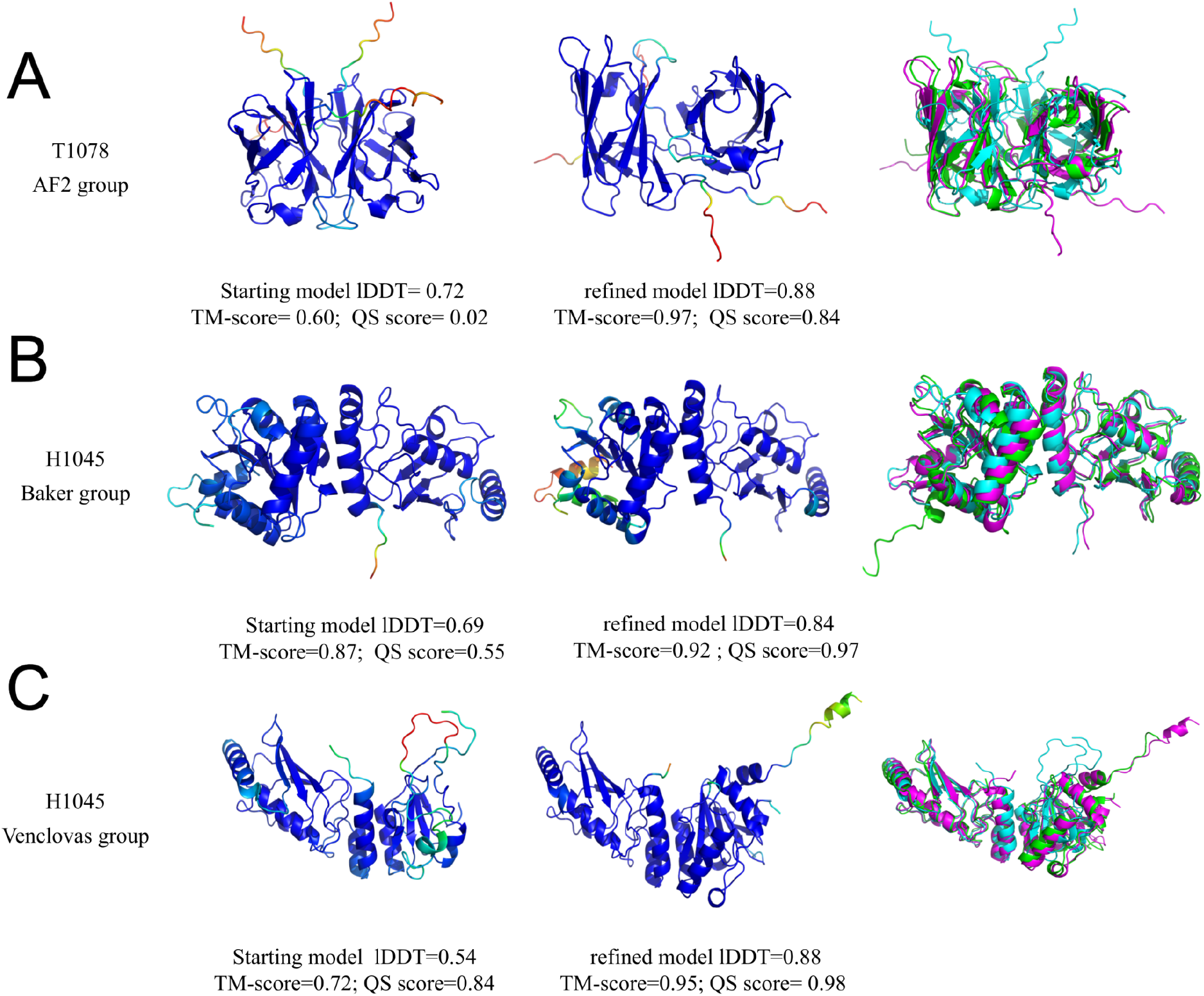
Comparison of multimeric models. Images in the left columns show the starting models coloured by plDDT score. The middle columns show the refined models coloured by plDDT score. The right columns show the superposition of starting models (cyan), the best-refined models generated by colabfold (magenta) and the observed models (green). (A) AF2 model for T1078: starting model lDDT=0.72, TM-score= 0.60, QS score=0.02, refined model lDDT=0.88, TM-score=0.97, QS score= 0.84. (B) Baker group model for H1045: starting model lDDT=0.69, TM-score= 0.87, QS score=0.55, refined lDDT=0.84, TM-score=0.92, QS score=0.97. (C) Venclovas group model for H1045: starting model lDDT=0.54,TM-score=0.72, QS score=0.84, refined lDDT=0.88, TM-score=0.95, QS score=0.98. Images were rendered using PyMOL.

The difference in performance between monomer and complex recycling experiments; namely that monomers show a greater improvement in quality when measured by the lDDT metric whereas complexes tend to show greater improvement when measured by TM-Score, is likely a consequence of the different focus of calibration used in the development of the two AF2 versions. AlphaFold2 is primarily calibrated to lDDT (Jumper et al., 2021; Tunyasuvunakool et al., 2021) whereas AlphaFold2-Multimer is calibrated to TM-Score (Evans et al., 2021).

## 4. Conclusions

We have demonstrated that ReFOLD4 is successful in its main goal of preventing MD simulations from structural deviations and is a risk-averse method for providing the conformational landscape of AF2 accuracy level predictions for further studies such as drug discovery. It is also promising that by generating a higher population of improved models on FM targets, ReFOLD4 may be of use in de novo protein design pipelines. Molprobity score analysis showed that 94 percent of models generated by ReFOLD4 were improved, and all models were improved for 46 targets out of 57 compared to the starting models generated by LocalColabFold. 72 percent of the models were also improved when the best AF2 submissions were used as starting models. It is also worthy of note that our refinement pipeline managed to provide better-improved models for 18 out of 57 targets compared to the AlphaFold group (427) according to the maximum GDT-TS score. The selection of ReFOLD4’s best-improved model remains a challenge, despite its high number of improved models.

Furthermore, we have demonstrated that the AF2 algorithm can be used to refine 3D models when they are input as custom templates. If the main aim is to consistently improve the quality of a single model, either mono- or multi-meric, then the AF2 recycling process is advantageous in terms of the relatively low computational resources required and it provides a ranking of models for easy selection by predicted quality score (e.g., plDDT or pTM). Both MSA and single sequence input recycling led to a significant improvement in model quality compared to baseline. Importantly, this improvement occurred not only with non-AF models but with original AF2 models that were submitted by DeepMind in CASP14. The lack of significant improvement in consecutive recycles showed that a higher recycle number did not necessarily lead to greater improvement. Comparing the p-values, recycle 3 appeared to improve model quality as much as any other recycle and therefore committing monomeric structures to more than 3 recycles may represent an unnecessary processing overhead. This is shown in Figure 3 by a change in cumulative observed lDDT score of 7.54 for recycle 3 compared to 6.34 at recycle 12 (AF2 models) and a modest 2.7% increase in cumulative observed lDDT score across all non-AF models from recycle 3 to 12. Lastly, for multimeric models, instead of a recycle value of 3, further recycling appears to be required and, as a general rule, recycle 12 in MSA mode (26% increase) was most successful as measured by IDDT and QS-score.

A combined approach to refinement using both the ReFOLD4 protocol and the AF2 recycling processes would provide a platform for further conformational studies, such as drug discovery. It also provides substantial improvements to 3D models beyond the AF2 accuracy level while requiring modest additional computational resources.

## Availability

Refinement using AlphaFold2-Multimer recycling can be carried out following our protocols using ColabFold (https://github.com/sokrypton/ColabFold) and is also part of the MultiFOLD pipline (https://hub.docker.com/r/mcguffin/multifold). ReFOLD4 scripts for MD using NAMD are available on request from the authors.

