## Supplementary for "Improvement of protein tertiary and quaternary structure predictions using the ReFOLD4 refinement method and the AlphaFold2 recycling process"

**Supplementary Table 1.** Performance summary for ReFOLD4 on the CASP14 targets according to Molprobity score (lower Molprobity scores are better). The starting 3D models generated by LocalColabFold were further refined by ReFOLD4.

| Target ID | Prediction Method | Score of best model submitted by AF2 in CASP14 | Score of starting model generated by ColabFold | Minimum score of refined models | Mean score of refined models | Maximum score of refined models | Percentage of improved models |
|---|---|---|---|---|---|---|---|
| T1027 | FM | 2.39 | 1.16 | 0.68 | 1.1432 | 1.58 | 63.414 |
| T1029 | FM | 0.79 | 1.74 | 0.5 | 0.883 | 1.41 | 100 |
| T1031 | FM | 1.62 | 1.41 | 0.54 | 0.985 | 1.58 | 98.170 |
| T1033 | FM | 1.51 | 1.55 | 0.51 | 0.920 | 1.39 | 100 |
| T1037 | FM | 0.9 | 1.33 | 0.73 | 1.080 | 1.32 | 100 |
| T1039 | FM | 1.84 | 0.88 | 0.5 | 0.982 | 1.34 | 27.439 |
| T1040 | FM | 0.5 | 1.75 | 0.5 | 0.842 | 1.36 | 100 |
| T1041 | FM | 1 | 1 | 0.52 | 0.937 | 1.28 | 73.780 |
| T1042 | FM | 1.35 | 3.66 | 0.76 | 1.042 | 1.33 | 100 |
| T1043 | FM | 1.2 | 1.28 | 0.52 | 1.002 | 1.52 | 96.341 |
| T1047s1 | FM | 1.61 | 1.35 | 0.77 | 1.101 | 1.48 | 93.292 |
| T1049 | FM | 0.56 | 0.89 | 0.8 | 1.157 | 1.58 | 12.195 |
| T1064 | FM | 1.14 | 1.51 | 0.5 | 1.125 | 1.57 | 98.170 |
| T1074 | FM | 1.14 | 1.84 | 0.97 | 1.229 | 1.81 | 100 |
| T1090 | FM | 0.68 | 0.5 | 0.5 | 0.899 | 1.38 | 0 |
| T1035 | FM/TBM | 1.02 | 2.35 | 0.5 | 0.764 | 1.32 | 100 |
| T1038 | FM/TBM | 0.83 | 2.56 | 0.5 | 0.972 | 1.8 | 100 |
| T1046s1 | FM/TBM | 1.03 | 2.3 | 0.5 | 0.771 | 1.3 | 100 |
| T1047s2 | FM/TBM | 0.79 | 2.47 | 0.71 | 1.021 | 1.26 | 100 |
| T1052 | FM/TBM | 1.07 | 2.26 | 0.78 | 1.161 | 1.38 | 100 |
| T1053 | FM/TBM | 1.01 | 2.1 | 0.71 | 1.117 | 1.34 | 100 |

| | | | | | | | |
|---|---|---|---|---|---|---|---|
| T1055 | FM/TBM | 1.19 | 2.3 | 0.5 | 0.7148 | 1.2 | 99.393 |
| T1058 | FM/TBM | 0.83 | 1.88 | 0.53 | 0.909 | 1.26 | 100 |
| T1061 | FM/TBM | 1.16 | 2.15 | 0.78 | 1.120 | 1.35 | 100 |
| T1065s2 | FM/TBM | 1.19 | 1.68 | 0.5 | 0.831 | 1.33 | 100 |
| T1070 | FM/TBM | 1.41 | 2.95 | 0.89 | 1.268 | 1.53 | 100 |
| T1082 | FM/TBM | 1.22 | 2.79 | 0.5 | 0.919 | 1.39 | 100 |
| T1093 | FM/TBM | 0.9 | 2.12 | 0.55 | 0.812 | 1.09 | 100 |
| T1094 | FM/TBM | 1.18 | 2.4 | 0.72 | 1.011 | 1.31 | 100 |
| T1024 | TBM | 1.06 | 2.17 | 0.5 | 0.795 | 1.11 | 100 |
| T1026 | TBM | 1.83 | 3.13 | 0.59 | 1.162 | 1.83 | 100 |
| T1030 | TBM | 0.72 | 1.63 | 0.5 | 1.188 | 0.97 | 100 |
| T1032 | TBM | 0.66 | 1.77 | 0.5 | 0.844 | 1.18 | 100 |
| T1034 | TBM | 1.43 | 2.23 | 0.5 | 0.939 | 1.31 | 100 |
| T1045s2 | TBM | 1.11 | 1.81 | 0.5 | 0.723 | 1.11 | 100 |
| T1046s2 | TBM | 1.56 | 2.08 | 0.53 | 1.00 | 1.56 | 100 |
| T1050 | TBM | 1.06 | 1.91 | 0.68 | 1.04 | 1.27 | 100 |
| T1054 | TBM | 1.53 | 2.67 | 0.9 | 1.195 | 1.57 | 100 |
| T1056 | TBM | 1.07 | 2.72 | 0.84 | 1.14 | 1.54 | 100 |
| T1060s2 | TBM | 1.18 | 2.27 | 0.5 | 1.018 | 1.52 | 100 |
| T1060s3 | TBM | 0.67 | 2.27 | 0.5 | 1.03 | 1.54 | 100 |
| T1065s1 | TBM | 0.5 | 1.96 | 0.5 | 0.811 | 1.38 | 100 |
| T1067 | TBM | 0.79 | 2.49 | 0.79 | 1.068 | 1.37 | 100 |
| T1068 | TBM | 1.1 | 2.4 | 0.58 | 0.958 | 1.34 | 100 |
| T1073 | TBM | 1.1 | 2.24 | 0.5 | 0.774 | 1.26 | 100 |
| T1076 | TBM | 0.88 | 1.78 | 0.5 | 1.0738 | 1.37 | 100 |
| T1078 | TBM | 1.43 | 2.38 | 0.5 | 0.989 | 1.41 | 100 |
| T1079 | TBM | 0.87 | 2.25 | 0.87 | 1.068 | 1.35 | 100 |

| | | | | | | | |
|---|---|---|---|---|---|---|---|
| T1083 | TBM | 0.71 | 1.78 | 0.5 | 0.704 | 1.22 | 100 |
| T1084 | TBM | 0.74 | 2.02 | 0.5 | 0.579 | 1.11 | 100 |
| T1087 | TBM | 0.99 | 2.65 | 0.5 | 0.935 | 1.55 | 100 |
| T1089 | TBM | 1.24 | 2.39 | 0.96 | 1.248 | 1.51 | 100 |
| T1092 | TBM | 0.88 | 2.17 | 0.58 | 0.893 | 1.17 | 100 |
| T1095 | TBM | 1.12 | 2.25 | 0.77 | 0.956 | 1.26 | 100 |
| T1099 | TBM | 1.49 | 2.96 | 0.76 | 1.119 | 1.45 | 100 |
| T1100 | TBM | 1.11 | 1.85 | 0.5 | 0.821 | 1.17 | 100 |
| T1101 | TBM | 1.02 | 2.39 | 0.63 | 1.030 | 1.4 | 100 |
| The Cumulative scores | | 62.91 | 116.78 | 34.95 | 55.86 | 78.62 | |

**Supplementary Table 2.** Performance summary for ReFOLD4 on the AF2 CASP14 FM models according to GDT-TS, and lDDT scores. The best models submitted by AlphaFold2 group (427) were further refined by ReFOLD4.

| | GDT-TS score | | | | | lDDT score | | | | |
|---|---|---|---|---|---|---|---|---|---|---|
| CASP Target ID | Score of starting model submitted by AF2 in CASP14 | Minimum score of refined models | Mean score of refined models | Maximum score of refined models | Percentage of improved models | Score of starting model submitted by AF2 in CASP14 | Minimum score of refined models | Mean score of refined models | Maximum score of refined models | Percentage of improved models |
| T1027 | 0.388 | 0.372 | 0.382 | 0.394 | 12.1951 | 0.433 | 0.413 | 0.418 | 0.431 | 0 |
| T1029 | 0.446 | 0.428 | 0.442 | 0.452 | 28.658 | 0.472 | 0.456 | 0.461 | 0.470 | 0 |
| T1031 | 0.871 | 0.828 | 0.856 | 0.878 | 7.317 | 0.714 | 0.671 | 0.687 | 0.711 | 0 |
| T1033 | 0.877 | 0.832 | 0.860 | 0.877 | 3.658 | 0.81 | 0.7513 | 0.773 | 0.809 | 0 |
| T1037 | 0.872 | 0.829 | 0.847 | 0.873 | 3.65 | 0.79 | 0.744 | 0.755 | 0.792 | 0 |
| T1039 | 0.823 | 0.788 | 0.805 | 0.818 | 0 | 0.71 | 0.675 | 0.691 | 0.714 | 0 |
| T1040 | 0.719 | 0.694 | 0.713 | 0.732 | 23.170 | 0.74 | 0.711 | 0.7303 | 0.742 | 4.8788 |

| | | | | | | | | | |
|---|---|---|---|---|---|---|---|---|---|
| T1041 | 0.905 | 0.862 | 0.883 | 0.9081 | 3.048 | 0.82 | 0.76 | 0.783 | 0.821 | 0 |
| T1042 | 0.837 | 0.789 | 0.808 | 0.838 | 1.215 | 0.83 | 0.77 | 0.775 | 0.829 | 0 |
| T1043 | 0.832 | 0.783 | 0.808 | 0.831 | 0 | 0.75 | 0.69 | 0.716 | 0.743 | 0 |
| T1047s1 | 0.502 | 0.477 | 0.489 | 0.5 | 0 | 0.75 | 0.714 | 0.725 | 0.745 | 0 |
| T1049 | 0.932 | 0.889 | 0.9107 | 0.931 | 0 | 0.843 | 0.780 | 0.795 | 0.840 | 0 |
| T1064 | 0.803 | 0.762 | 0.789 | 0.811 | 6.0976 | 0.746 | 0.673 | 0.688 | 0.733 | 0 |
| T1074 | 0.899 | 0.859 | 0.878 | 0.901 | 3.04878 | 0.836 | 0.751 | 0.770 | 0.832 | 0 |
| T1090 | 0.888 | 0.863 | 0.874 | 0.890 | 1.2195 | 0.827 | 0.778 | 0.791 | 0.824 | 0 |
| The Cumulative scores | 11.601 | 11.063 | 11.3524 | 11.638 | | 11.108 | 10.362 | 10.56 | 11.0449 | |

**Supplementary Table 3.** Performance summary for ReFOLD4 on the CASP14 targets according to GDT-TS score. The starting 3D models generated by LocalColabFold were further refined by ReFOLD4.

| CASP Target ID | Prediction Method | Score of best model submitted by AF2 in CASP14 | Score of starting model generated by ColabFold | Minimum score of refined models | Mean score of refined models | Maximum score of refined models | Percentage of improved models |
|---|---|---|---|---|---|---|---|
| T1027 | FM | 0.388 | 0.382 | 0.358 | 0.368 | 0.383 | 4.242 |
| T1029 | FM | 0.446 | 0.438 | 0.42 | 0.434 | 0.446 | 29.878 |
| T1031 | FM | 0.871 | 0.844 | 0.807 | 0.827 | 0.844 | 3.0487 |
| T1033 | FM | 0.877 | 0.845 | 0.812 | 0.834 | 0.857 | 17.073 |
| T1037 | FM | 0.872 | 0.729 | 0.703 | 0.718 | 0.729 | 1.219 |
| T1039 | FM | 0.823 | 0.843 | 0.796 | 0.8168 | 0.835 | 0 |
| T1040 | FM | 0.719 | 0.548 | 0.532 | 0.546 | 0.557 | 45.731 |
| T1041 | FM | 0.905 | 0.874 | 0.848 | 0.865 | 0.879 | 11.585 |

| | | | | | | | |
|---|---|---|---|---|---|---|---|
| T1042 | FM | 0.837 | 0.633 | 0.608 | 0.621 | 0.635 | 4.268 |
| T1043 | FM | 0.832 | 0.785 | 0.758 | 0.779 | 0.795 | 25.609 |
| T1047s1 | FM | 0.502 | 0.498 | 0.484 | 0.494 | 0.504 | 17.682 |
| T1049 | FM | 0.932 | 0.944 | 0.906 | 0.925 | 0.942 | 0 |
| T1064 | FM | 0.803 | 0.573 | 0.541 | 0.562 | 0.583 | 10.975 |
| T1074 | FM | 0.899 | 0.929 | 0.884 | 0.901 | 0.922 | 0 |
| T1090 | FM | 0.888 | 0.883 | 0.854 | 0.871 | 0.884 | 2.439 |
| T1035 | FM/TBM | 0.953 | 0.887 | 0.845 | 0.875 | 0.894 | 9.756 |
| T1038 | FM/TBM | 0.868 | 0.873 | 0.831 | 0.851 | 0.876 | 1.829 |
| T1046s1 | FM/TBM | 0.975 | 0.968 | 0.944 | 0.963 | 0.975 | 27.439 |
| T1047s2 | FM/TBM | 0.669 | 0.720 | 0.692 | 0.706 | 0.724 | 3.658 |
| T1052 | FM/TBM | 0.582 | 0.561 | 0.548 | 0.555 | 0.563 | 4.878 |
| T1053 | FM/TBM | 0.893 | 0.937 | 0.9 | 0.916 | 0.938 | 1.829 |
| T1055 | FM/TBM | 0.864 | 0.873 | 0.813 | 0.836 | 0.870 | 0 |
| T1058 | FM/TBM | 0.858 | 0.836 | 0.8 | 0.820 | 0.839 | 5.487 |
| T1061 | FM/TBM | 0.615 | 0.633 | 0.620 | 0.629 | 0.638 | 15.853 |
| T1065s2 | FM/TBM | 0.989 | 0.977 | 0.933 | 0.958 | 0.974 | 0 |
| T1070 | FM/TBM | 0.410 | 0.415 | 0.400 | 0.407 | 0.416 | 3.048 |
| T1082 | FM/TBM | 0.953 | 0.976 | 0.92 | 0.960 | 0.98 | 5.48 |
| T1093 | FM/TBM | 0.677 | 0.509 | 0.506 | 0.526 | 0.539 | 97.575 |
| T1094 | FM/TBM | 0.707 | 0.673 | 0.667 | 0.680 | 0.693 | 92.682 |
| T1024 | TBM | 0.600 | 0.593 | 0.571 | 0.582 | 0.593 | 0.609 |
| T1026 | TBM | 0.934 | 0.885 | 0.811 | 0.834 | 0.883 | 0 |
| T1030 | TBM | 0.628 | 0.636 | 0.609 | 0.622 | 0.637 | 1.2195 |
| T1032 | TBM | 0.686 | 0.682 | 0.663 | 0.676 | 0.685 | 14.634 |

| | | | | | | | |
|---|---|---|---|---|---|---|---|
| T1034 | TBM | 0.937 | 0.943 | 0.921 | 0.939 | 0.950 | 24.390 |
| T1045s2 | TBM | 0.917 | 0.917 | 0.875 | 0.899 | 0.921 | 3.0487 |
| T1046s2 | TBM | 0.964 | 0.957 | 0.934 | 0.951 | 0.966 | 21.951 |
| T1050 | TBM | 0.860 | 0.799 | 0.774 | 0.786 | 0.801 | 3.658 |
| T1054 | TBM | 0.921 | 0.898 | 0.867 | 0.883 | 0.898 | 3.658 |
| T1056 | TBM | 0.952 | 0.849 | 0.825 | 0.835 | 0.849 | 1.829 |
| T1060s2 | TBM | 0.827 | 0.774 | 0.753 | 0.766 | 0.777 | 9.756 |
| T1060s3 | TBM | 0.951 | 0.920 | 0.900 | 0.925 | 0.943 | 78.048 |
| T1065s1 | TBM | 0.958 | 0.960 | 0.924 | 0.945 | 0.958 | 0 |
| T1067 | TBM | 0.893 | 0.895 | 0.855 | 0.879 | 0.896 | 3.048 |
| T1068 | TBM | 0.962 | 0.967 | 0.946 | 0.958 | 0.969 | 2.439 |
| T1073 | TBM | 0.839 | 0.826 | 0.783 | 0.808 | 0.826 | 2.439 |
| T1076 | TBM | 0.990 | 0.986 | 0.950 | 0.963 | 0.986 | 3.048 |
| T1078 | TBM | 0.959 | 0.967 | 0.949 | 0.963 | 0.970 | 28.048 |
| T1079 | TBM | 0.916 | 0.913 | 0.880 | 0.896 | 0.913 | 1.219 |
| T1083 | TBM | 0.856 | 0.877 | 0.847 | 0.864 | 0.883 | 7.317 |
| T1084 | TBM | 0.912 | 0.915 | 0.887 | 0.905 | 0.926 | 17.682 |
| T1087 | TBM | 0.967 | 0.825 | 0.79 | 0.813 | 0.828 | 5.487 |
| T1089 | TBM | 0.970 | 0.974 | 0.952 | 0.963 | 0.976 | 3.655 |
| T1092 | TBM | 0.738 | 0.495 | 0.49 | 0.505 | 0.512 | 98.780 |
| T1095 | TBM | 0.704 | 0.7106 | 0.713 | 0.726 | 0.736 | 100 |
| T1099 | TBM | 0.751 | 0.695 | 0.668 | 0.684 | 0.700 | 4.268 |
| T1100 | TBM | 0.792 | 0.797 | 0.767 | 0.784 | 0.802 | 6.097 |
| T1101 | TBM | 0.870 | 0.824 | 0.791 | 0.809 | 0.827 | 7.926 |

| The Cumulative scores | | 46.861 | 45.0656 | 43.425 | 44.4068 | 45.325 |
|---|---|---|---|---|---|---|

**Supplementary Table 4**. Performance summary for ReFOLD4 on the CASP14 targets according to lDDTscore. The starting 3D models generated by LocalColabFold were further refined by ReFOLD4

| CASP Target ID | Prediction method | Score of model submitted by AF2 in CASP14 | The starting model generated by ColabFold | Minimum score of refined models | Mean score of refined models | Maximum score of refined models | Percentage of improved models |
|---|---|---|---|---|---|---|---|
| T1027 | FM | 0.433 | 0.464 | 0.433 | 0.441 | 0.464 | 0.609 |
| T1029 | FM | 0.472 | 0.472 | 0.456 | 0.461 | 0.470 | 0 |
| T1031 | FM | 0.714 | 0.696 | 0.665 | 0.681 | 0.704 | 4.268 |
| T1033 | FM | 0.818 | 0.757 | 0.709 | 0.722 | 0.749 | 0 |
| T1037 | FM | 0.792 | 0.703 | 0.670 | 0.678 | 0.704 | 3.0487 |
| T1039 | FM | 0.717 | 0.734 | 0.689 | 0.703 | 0.728 | 0 |
| T1040 | FM | 0.740 | 0.635 | 0.605 | 0.616 | 0.633 | 0 |
| T1041 | FM | 0.828 | 0.834 | 0.78 | 0.798 | 0.834 | 0 |
| T1042 | FM | 0.831 | 0.673 | 0.635 | 0.645 | 0.673 | 2.439 |
| T1043 | FM | 0.75 | 0.749 | 0.702 | 0.723 | 0.743 | 0 |
| T1047s1 | FM | 0.75 | 0.745 | 0.712 | 0.722 | 0.746 | 3.0487 |
| T1049 | FM | 0.848 | 0.868 | 0.792 | 0.810 | 0.86 | 0 |
| T1064 | FM | 0.746 | 0.515 | 0.461 | 0.478 | 0.508 | 0 |
| T1074 | FM | 0.836 | 0.846 | 0.761 | 0.776 | 0.836 | 0 |
| T1090 | FM | 0.827 | 0.830 | 0.782 | 0.793 | 0.828 | 0 |
| T1035 | FM/TBM | 0.868 | 0.824 | 0.757 | 0.781 | 0.813 | 0 |

| | | | | | | | |
|---|---|---|---|---|---|---|---|
| T1038 | FM/TBM | 0.832 | 0.830 | 0.777 | 0.789 | 0.825 | 0 |
| T1046s1 | FM/TBM | 0.894 | 0.882 | 0.816 | 0.837 | 0.88 | 0 |
| T1047s2 | FM/TBM | 0.775 | 0.788 | 0.747 | 0.758 | 0.787 | 0 |
| T1052 | FM/TBM | 0.864 | 0.853 | 0.800 | 0.810 | 0.854 | 0 |
| T1053 | FM/TBM | 0.860 | 0.853 | 0.806 | 0.815 | 0.851 | 0 |
| T1055 | FM/TBM | 0.737 | 0.746 | 0.649 | 0.680 | 0.730 | 0 |
| T1058 | FM/TBM | 0.826 | 0.813 | 0.765 | 0.775 | 0.811 | 0 |
| T1061 | FM/TBM | 0.728 | 0.733 | 0.710 | 0.716 | 0.734 | 3.048 |
| T1065s2 | FM/TBM | 0.909 | 0.913 | 0.818 | 0.849 | 0.905 | 0 |
| T1070 | FM/TBM | 0.730 | 0.734 | 0.691 | 0.701 | 0.735 | 2.439 |
| T1082 | FM/TBM | 0.866 | 0.870 | 0.793 | 0.820 | 0.857 | 0 |
| T1093 | FM/TBM | 0.749 | 0.737 | 0.730 | 0.735 | 0.741 | 9.756 |
| T1094 | FM/TBM | 0.754 | 0.745 | 0.732 | 0.738 | 0.743 | 0 |
| T1024 | TBM | 0.781 | 0.772 | 0.733 | 0.743 | 0.772 | 2.439 |
| T1026 | TBM | 0.808 | 0.723 | 0.658 | 0.672 | 0.721 | 0 |
| T1030 | TBM | 0.850 | 0.876 | 0.812 | 0.829 | 0.873 | 0 |
| T1032 | TBM | 0.72 | 0.723 | 0.676 | 0.688 | 0.722 | 0 |
| T1034 | TBM | 0.853 | 0.864 | 0.813 | 0.826 | 0.863 | 0 |
| T1045s2 | TBM | 0.852 | 0.853 | 0.792 | 0.810 | 0.851 | 0 |
| T1046s2 | TBM | 0.877 | 0.875 | 0.815 | 0.831 | 0.874 | 0 |
| T1050 | TBM | 0.871 | 0.868 | 0.819 | 0.828 | 0.869 | 3.658 |
| T1054 | TBM | 0.869 | 0.857 | 0.790 | 0.812 | 0.856 | 0 |
| T1056 | TBM | 0.904 | 0.812 | 0.740 | 0.757 | 0.811 | 0 |
| T1060s2 | TBM | 0.897 | 0.897 | 0.833 | 0.856 | 0.892 | 0 |
| T1060s3 | TBM | 0.815 | 0.804 | 0.780 | 0.794 | 0.806 | 3.048 |
| T1065s1 | TBM | 0.897 | 0.897 | 0.833 | 0.856 | 0.892 | 0 |
| T1067 | TBM | 0.869 | 0.866 | 0.802 | 0.820 | 0.865 | 0 |

| | | | | | | | |
|---|---|---|---|---|---|---|---|
| T1068 | TBM | 0.905 | 0.908 | 0.834 | 0.852 | 0.899 | 0 |
| T1073 | TBM | 0.771 | 0.768 | 0.680 | 0.707 | 0.756 | 0 |
| T1076 | TBM | 0.943 | 0.937 | 0.864 | 0.875 | 0.935 | 0 |
| T1078 | TBM | 0.922 | 0.941 | 0.857 | 0.880 | 0.936 | 0 |
| T1079 | TBM | 0.909 | 0.915 | 0.861 | 0.870 | 0.915 | 1.219 |
| T1083 | TBM | 0.782 | 0.817 | 0.753 | 0.774 | 0.812 | 0 |
| T1084 | TBM | 0.892 | 0.864 | 0.804 | 0.827 | 0.8613 | 0 |
| T1087 | TBM | 0.9 | 0.734 | 0.680 | 0.696 | 0.73 | 0 |
| T1089 | TBM | 0.917 | 0.924 | 0.864 | 0.877 | 0.9217 | 0 |
| T1092 | TBM | 0.766 | 0.775 | 0.773 | 0.777 | 0.7844 | 89.634146 |
| T1095 | TBM | 0.752 | 0.749 | 0.748 | 0.752 | 0.758 | 98.17073 |
| T1099 | TBM | 0.779 | 0.749 | 0.710 | 0.723 | 0.7501 | 3.0487804 |
| T1100 | TBM | 0.865 | 0.852 | 0.805 | 0.8184 | 0.8524 | 1.219512 |
| T1101 | TBM | 0.859 | 0.856 | 0.787 | 0.801 | 0.853 | 0 |
| The Cumulative scores | | 46.342 | 45.347 | 42.387 | 43.227 | 45.1978 | |

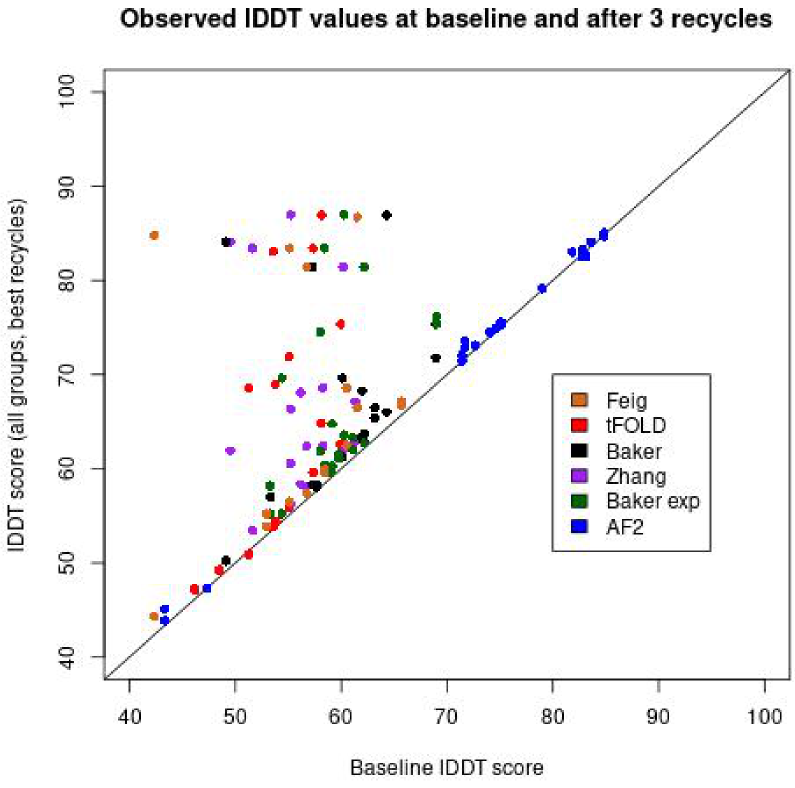

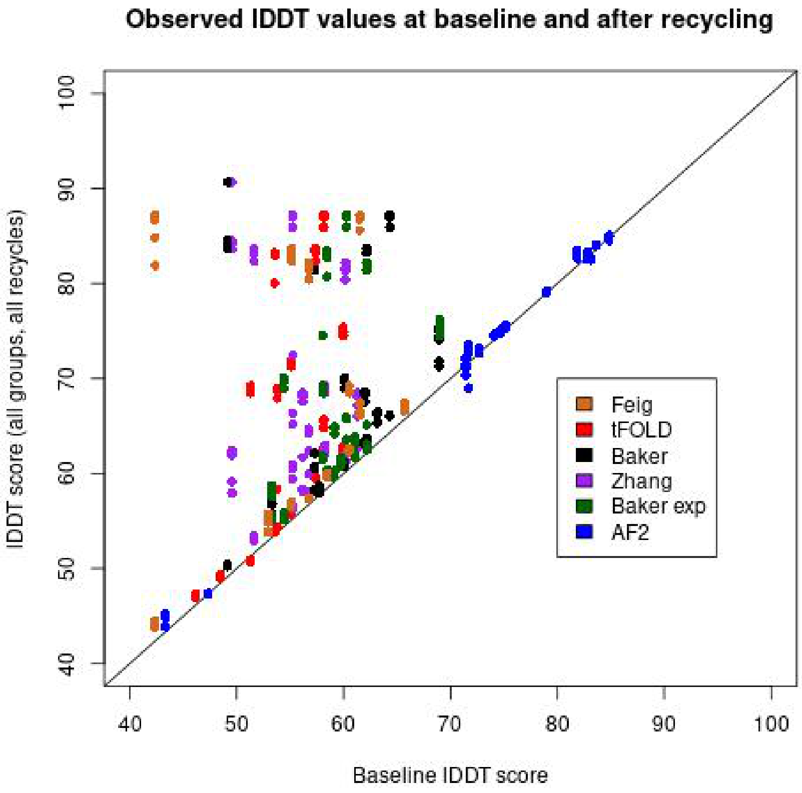

**Supplementary Figure 1.** Scatter plots showing observed lDDT scores for all models between baseline and recycle 3 (left) and baseline and all recycles (right) , coloured by group.

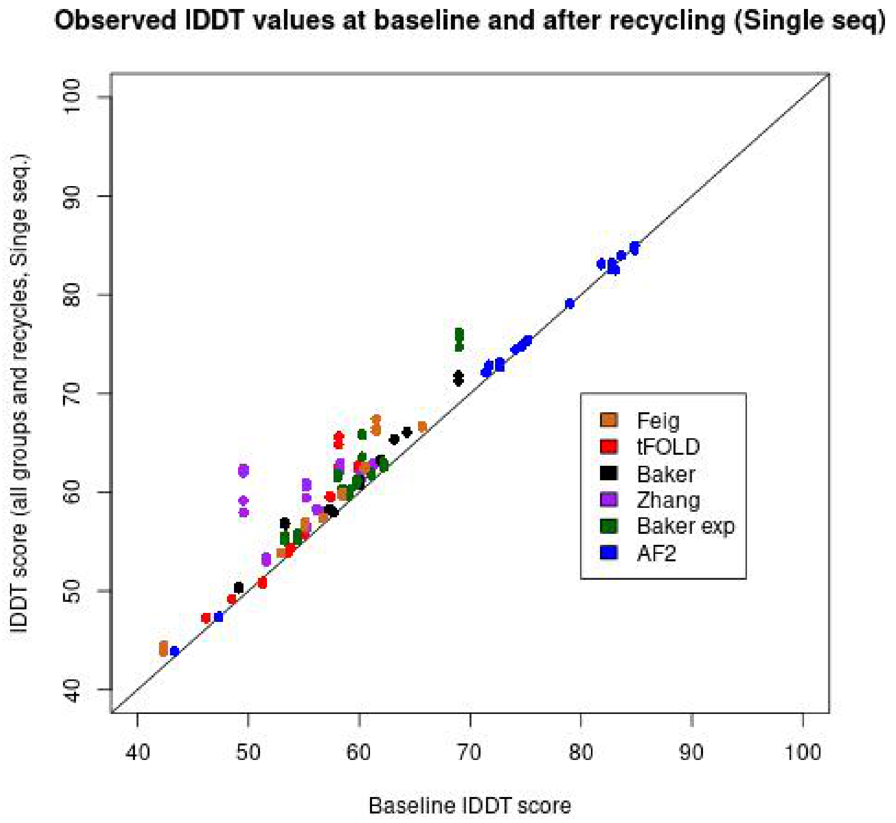

**Supplementary Figure 2:** Scatter plot to show comparisons in observed lDDT scores between baseline and all recycles for all models (AF2 and non-AF2).

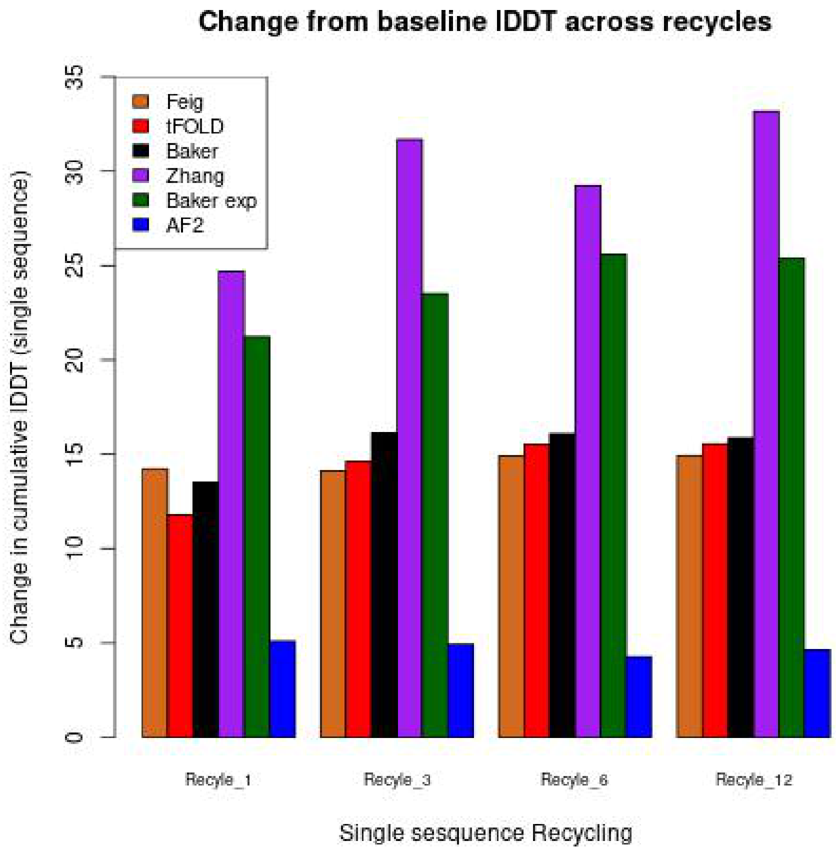

**Supplementary Figure 3:** Graphs to show average change in lDDT scores from baseline over all recycles for the individual CASP14 and AF2 models- single sequence modelling.

**Supplementary Table 5.** A comparison matrix of single-sequence versus MSA recycling by comparing mean lDDT scores and scale parameters across 1, 3, 6 and 12 recycles for CASP14 AF2 and non-AF2 models.

| | p-values (single sequence vs. MSA) | | | |
|---|---|---|---|---|
| **AF2 models** | Recycle 1 | Recycle 3 | Recycle 6 | Recycle 12 |
| Wilcox signed rank | 0.097 | 0.052 | 0.111 | 0.129 |
| Ansari test | 0.397 | 0.500 | 0.425 | 0.544 |
| **non-AF2 models** | p-values (single sequence vs. MSA) | | | |
| Wilcox signed rank | **$1.42 \times 10^{-14}$** | **$5.34 \times 10^{-9}$** | **$2.94 \times 10^{-12}$** | **$7.80 \times 10^{-9}$** |
| Ansari test | **0.014** | **0.015** | **0.019** | **0.012** |

Ho: Recycling using the single sequence setting produces models that are equal in quality to those produced using the MSA setting for equivalent recycle numbers. H1: Recycling using the single sequence setting produces models that are lower in quality to those produced using the MSA setting for equivalent recycle numbers. P-values ≤0.05 indicate significant differences. The 1-tailed Wilcoxon signed-rank sum test and 1-tailed Ansari-Bradley test were used to calculate p-values from lDDT scores across 16 AlphaFold CASP14 top-ranked models (upper two rows) and 47 non-AlphaFold CASP14 top-ranked models (lower two rows).

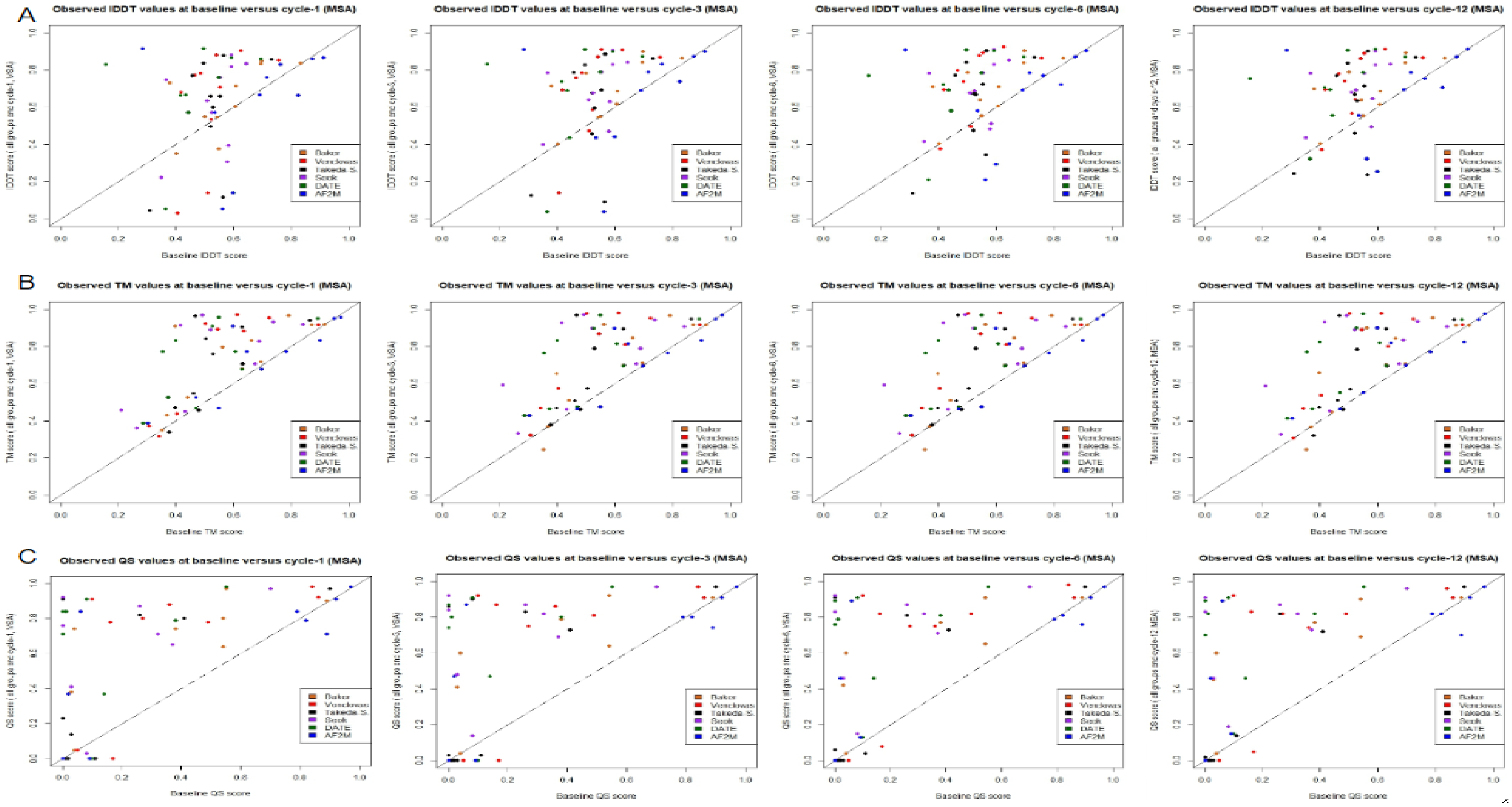

**Supplementary Figure 4:** Graphs to show comparisons in observed IDDT scores (A), TM scores (B), QS-scores (C) between baseline and each recycle separately for all models (AF2 and non-AF2) in the MSA methods.

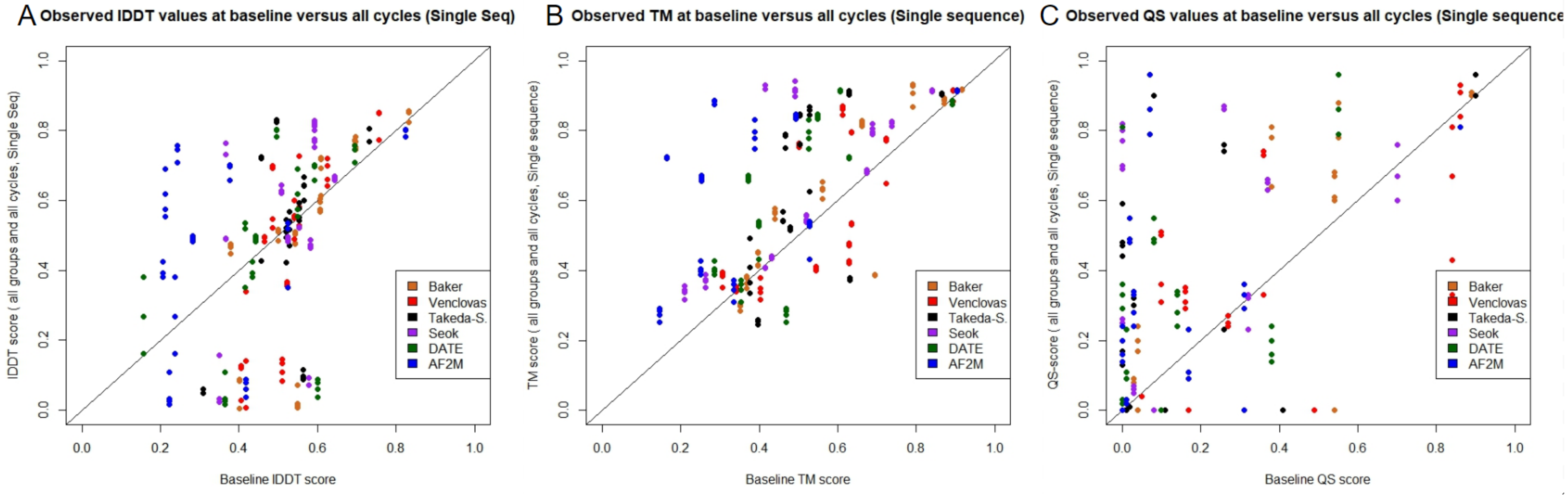

**Supplementary Figure 5 :** Graphs to show comparisons in observed lDDT scores (A), TM scores (B), QS-scores (C) between baseline and all recycle for all models (AF2 and non-AF2).

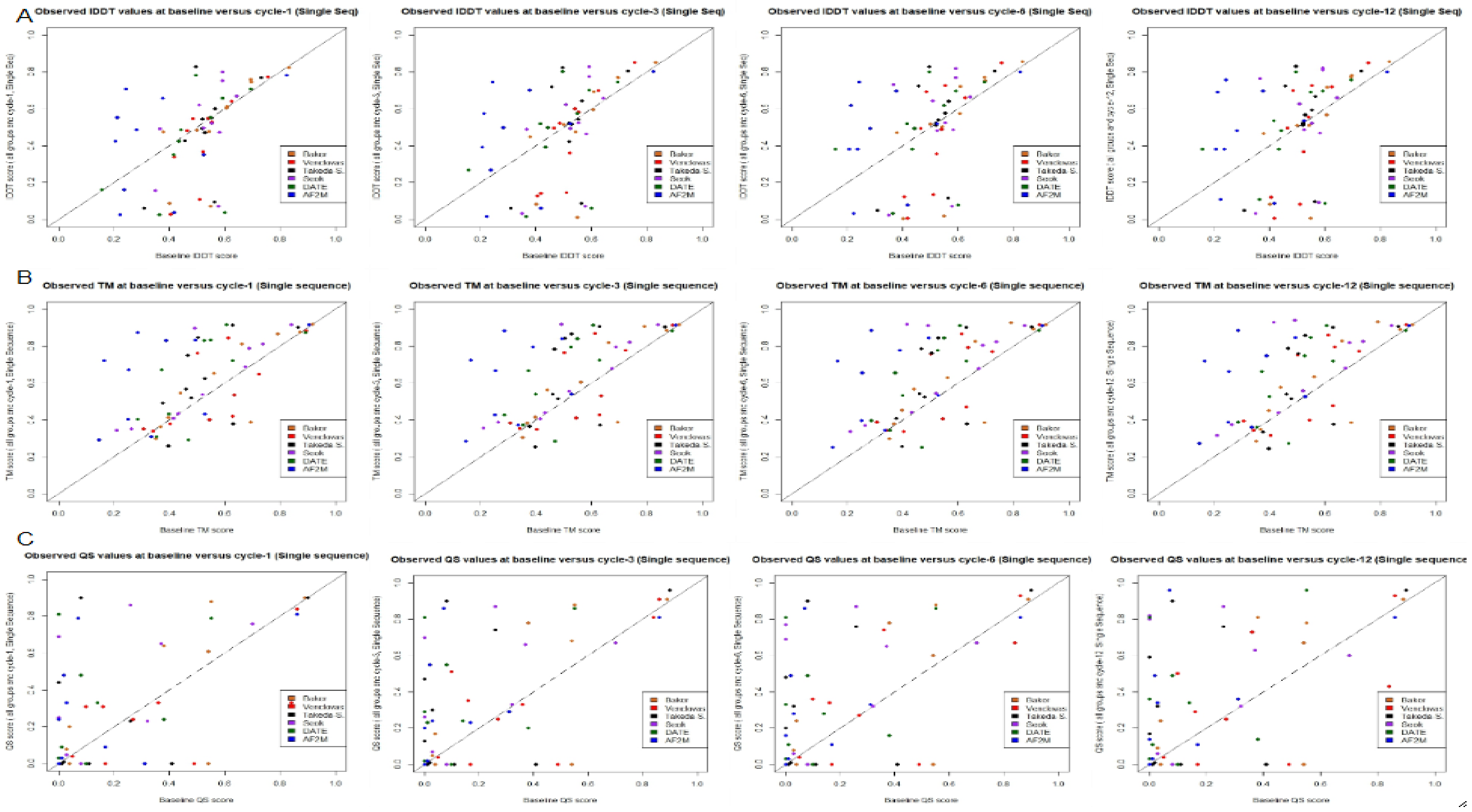

**Supplementary Figure 6 :** Graphs to show comparisons observed lDDT scores (A), TM scores (B), QS-scores (C) between baseline and each recycle separately, for all models (AF2 and non-AF2) in the Single Seq.
